# Auditory confounds can drive online effects of transcranial ultrasonic stimulation in humans

**DOI:** 10.1101/2023.02.22.527901

**Authors:** Benjamin R. Kop, Yazan Shamli Oghli, Talyta C. Grippe, Tulika Nandi, Judith Lefkes, Sjoerd W. Meijer, Soha Farboud, Marwan Engels, Michelle Hamani, Melissa Null, Angela Radetz, Umair Hassan, Ghazaleh Darmani, Andrey Chetverikov, Hanneke E.M. den Ouden, Til Ole Bergmann, Robert Chen, Lennart Verhagen

## Abstract

Transcranial ultrasonic stimulation (TUS) is rapidly emerging as a promising non-invasive neuromodulation technique. TUS is already well-established in animal models, providing foundations to now optimize neuromodulatory efficacy for human applications. Across multiple studies, one promising protocol, pulsed at 1000 Hz, has consistently resulted in motor cortical inhibition in humans (Fomenko et al., 2020). At the same time, a parallel research line has highlighted the potentially confounding influence of peripheral auditory stimulation arising from TUS pulsing at audible frequencies. In this study, we disentangle direct neuromodulatory and indirect auditory contributions to motor inhibitory effects of TUS. To this end, we include tightly matched control conditions across four experiments, one preregistered, conducted independently at three institutions. We employed a combined transcranial ultrasonic and magnetic stimulation paradigm, where TMS-elicited motor-evoked potentials (MEPs) served as an index of corticospinal excitability. First, we replicated motor inhibitory effects of TUS but showed through both tight controls and manipulation of stimulation intensity, duration, and auditory masking conditions that this inhibition was driven by peripheral auditory stimulation, not direct neuromodulation. Further, we consider neuromodulation beyond driving overall excitation/inhibition and show preliminary evidence of how TUS might interact with ongoing neural dynamics instead. Primarily, this study highlights the substantial shortcomings in accounting for the auditory confound in prior TUS-TMS work where only a flip-over sham and no active control was used. The field must critically reevaluate previous findings given the demonstrated impact of peripheral confounds. Further, rigorous experimental design via (in)active control conditions is required to make substantiated claims in future TUS studies. Only when direct effects are disentangled from those driven by peripheral confounds can TUS fully realize its potential for research and clinical applications.

## 1. Introduction

Noninvasive neuromodulation is a powerful tool for causal inference that strengthens our understanding of the brain and holds great clinical potential (Bergmann & Hartwigsen, 2021; Bestmann & Walsh, 2017). Transcranial ultrasonic stimulation (TUS) is a particularly promising non-invasive brain stimulation technique, overcoming current limitations with high spatial resolution and depth range (Darmani et al., 2022). The efficacy of TUS is well-established in cell cultures and animal models (Menz et al., 2013; Mohammadjavadi et al., 2019; Murphy et al., 2022; Tyler et al., 2008, 2018; Yoo et al., 2022), and emerging evidence for the neuromodulatory utility of TUS in humans has been reported for both cortical and subcortical structures (cortical: Butler et al., 2022; Lee et al., 2016; Liu et al., 2021; Zeng et al., 2022; subcortical: Ai et al., 2016; Cain et al., 2021; Nakajima et al., 2022). Especially now, at this foundational stage of TUS in humans, it is essential to converge on protocols that maximize the specificity and efficacy of stimulation (Folloni et al., 2019; Verhagen et al., 2019).

Motor inhibitory effects of a commonly applied 1000 Hz pulsed TUS protocol are among the most robust and replicable human findings (Fomenko et al., 2020; Legon, Bansal, et al., 2018; Xia et al., 2021). Here, by concurrently applying transcranial magnetic stimulation (TMS), modulation of corticospinal excitability is indexed by motor-evoked potentials (MEPs). However, the mechanism by which TUS evokes motor inhibition has remained under debate (Xia et al., 2021).

Recent studies in both animal and human models demonstrate how electrophysiological and behavioral outcomes of TUS can be elicited by nonspecific auditory activation rather than direct neuromodulation (Airan & Butts Pauly, 2018; Braun et al., 2020; Guo et al., 2018; Sato et al., 2018). Indeed, there is longstanding knowledge of the auditory confound accompanying pulsed TUS (Gavrilov & Tsirulnikov, 2012). However, this confound has only recently garnered attention, prompted by a pair of rodent studies demonstrating indirect auditory activation induced by TUS (Guo et al., 2022; Sato et al., 2018). Similar effects have been observed in humans, where exclusively auditory effects were captured with EEG measures (Braun et al., 2020). These findings are particularly impactful given that nearly all TUS studies employ pulsed protocols, from which the pervasive auditory confound emerges (Johnstone et al., 2021).

Indirect effects of stimulation are not unique to TUS, as transcranial magnetic and electric stimulation are also associated with auditory and somatosensory confounds. Indeed, the field of non-invasive brain stimulation as a whole depends on controlling for these confounding factors when present, to unveil the specificity of the neuromodulatory effects (Conde et al., 2019; Duecker et al., 2013; Polanía et al., 2018; Siebner et al., 2022). However, prior online TUS-TMS studies, including those exploring optimal neuromodulatory parameters to inform future work, have considered some but not all necessary conditions to control for the salient auditory confound elicited by a 1000 Hz pulsed protocol (Fomenko et al., 2020; Legon, Bansal, et al., 2018; Xia et al., 2021).

In this multicenter study, we quantified the impact of the auditory confound to disentangle direct neuromodulatory and indirect auditory contributions to motor inhibitory effects of TUS. To this end, we substantially improved upon prior TUS-TMS studies implementing solely flip-over sham by including both (in)active controls and multiple sound-sham conditions. Further, we investigated dose-response effects through administration of multiple stimulus durations, stimulation intensities, and individualized simulations of intracranial intensity. Additionally, we considered the possibility that online TUS might not drive a global change in the excitation/inhibition balance but instead might interact with ongoing neural dynamics by introducing state-dependent noise. Finally, we interrogated sound-driven effects through modulation of auditory confound volume, duration, pitch, and auditory masking. We show that motor inhibitory effects of TUS are spatially nonspecific and driven by sound-cued preparatory motor inhibition. However, we do find preliminary evidence that TUS might introduce dose– and state-dependent neural noise to the dynamics of corticospinal excitability. The present study highlights the importance of carefully constructed control conditions to account for confounding factors while exploring and refining TUS as a promising technique for human neuromodulation.

## 2. Materials and methods

### 2.1. Participants

This multicenter study comprised of four experiments conducted independently across three institutions. Experiment I (N = 12, 4 female, M_age_ = 25.9, SD_age_ = 4.6; METC: NL76920.091.21) and Experiment II (N = 27, 13 female, M_age_ = 24.1, SD_age_ = 3.7; METC: NL76920.091.21) were conducted at the Donders Institute of the Radboud University (the Netherlands). Experiment III was conducted at the Krembil Research Institute (N = 16, 8 female, M_age_ = 31.4, SD_age_ = 7.9; Toronto University Health Network Research Ethics Board: 20-5740, Canada), and Experiment IV at the Neuroimaging Centre of the Johannes Gutenberg University Medical Centre Mainz (N = 12, 11 female, M_age_ = 23.0, SD_age_ = 2.7, Landesärztekammer Rheinland-Pfalz: 2021-15808_01, Germany). All participants were healthy, right-handed, without a history of psychiatric or neurological disorders, and provided informed consent. Ethical approval was obtained for each experiment.

### 2.2. Transcranial ultrasonic and magnetic stimulation

Ultrasonic stimulation was delivered with the NeuroFUS system (manufacturer: Sonic Concepts Inc., Bothell, WA, USA; supplier/support: Brainbox Ltd., Cardiff, UK). A radiofrequency amplifier powered a piezoelectric ultrasound transducer via a matching network. Transducers consisted of a two-element annular array. Further transducer specifications are reported in **Supplementary Table 1**. Ultrasonic stimulation parameters were based on those used in prior TUS-TMS studies (**Table 1**, **Fig. 1A**; Fomenko et al., 2020; Legon, Bansal, et al., 2018; Xia et al., 2021). While ramping the pulses can in principle mitigate the auditory confound (Johnstone et al., 2021; Mohammadjavadi et al., 2019), doing so for such short pulse durations (<= 0.3 ms) is not effective. Therefore, we used a rectangular pulse shape to match prior work.

**Fig. 1.**
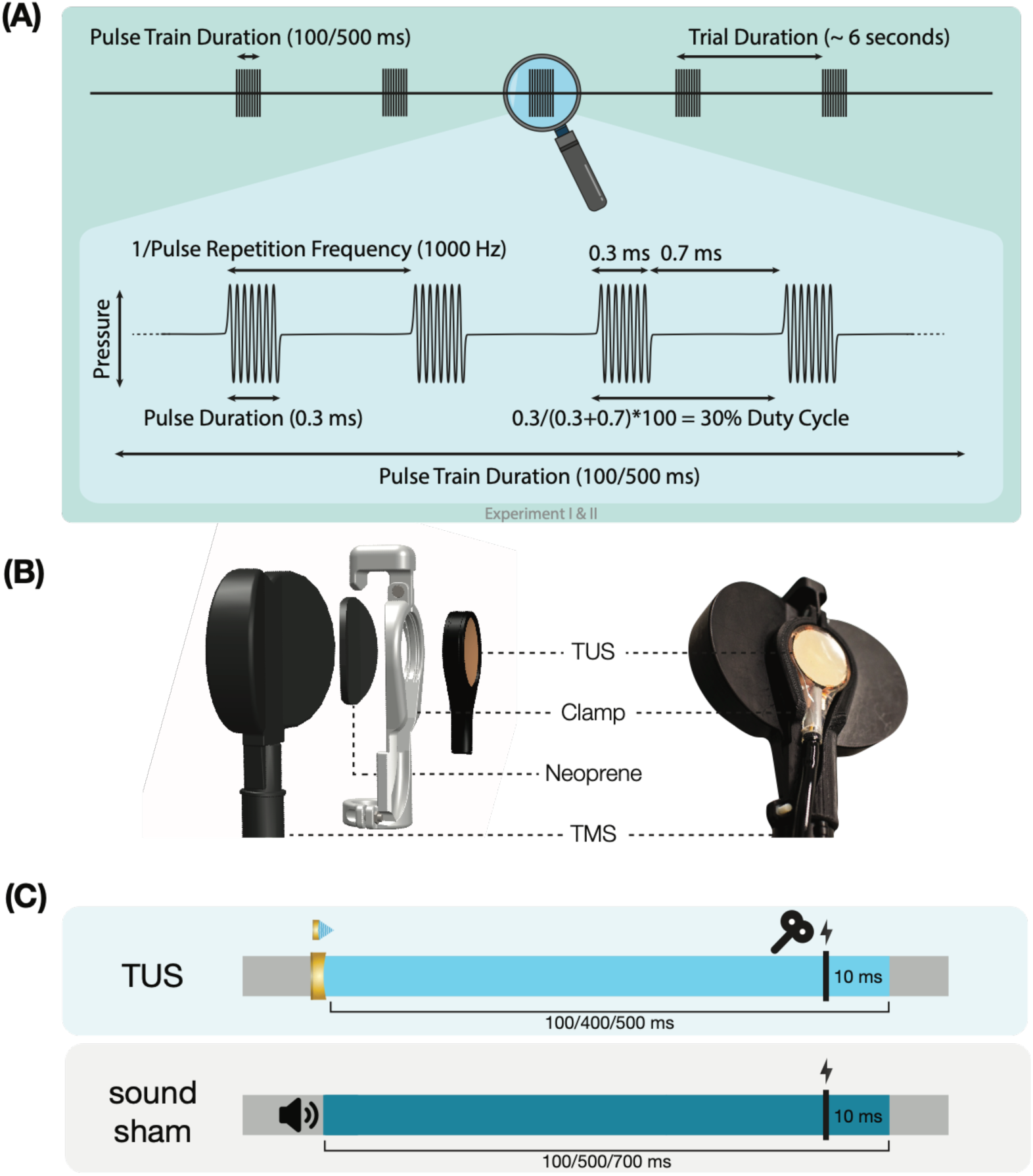
| Experimental procedures. **(A)** Ultrasonic stimulation protocol for Experiments I & II. In Experiment III a duty cycle of 10% was used. In Experiment IV a stimulus duration of 400 ms was used. **(B)** TUS-TMS clamp (<u>DOI:</u> <u>10.5281/zenodo.6517599</u>). **(C)** Experimental timing. Detailed experimental timing for each experiment is reported in **Supplementary** Fig. 3.

**Table 1.** | Ultrasonic stimulation parameters. *f* = fundamental frequency, depth = TPO focus setting for distance of free-water full-width half-maximum from transducer exit plane, PD = pulse duration, PRF = pulse repetition frequency, DC = duty cycle, PTD = pulse train duration, I_sppa_ = spatial-peak pulse-average intensity in free-water, P = pressure, MItc = transcranial derated mechanical index. The ramp shape for all experiments was rectangular. For estimated intracranial indices for Experiments I & II see **Supplementary** Figure 2.

| Exp. | $F$<br>(kHz) | depth<br>(mm) | PD<br>(ms) | PRF<br>(kHz) | DC | PTD<br>(ms) | $I_{sppa}$<br>(W/cm<sup>2</sup>) | P<br>(Mpa) | $MI_{tc}$ |
| --- | --- | --- | --- | --- | --- | --- | --- | --- | --- |
| I | 500 | 35 | 0.3 | 1000 | 30% | 100/500 | 32.5/65 | 1.02/1.44 | 0.99/1.40 |
| II | 250 | 28 | 0.3 | 1000 | 30% | 500 | 6.35/19.06 | 0.45/0.78 | 0.65/1.12 |
| III | 500 | 30 | 0.1 | 1000 | 10% | 500 | 9.26 | 0.54 | 0.53 |
| IV | 250 | 50<sup>†</sup> | 0.3 | 1000 | 30% | 400 | 4.34/8.69/10.52 | 0.37/0.53/0.58 | 0.53/0.76/0.83 |

<sup>†</sup>Note: Experiments I-III targeted the hand motor area. Experiment IV targeted the corticospinal white matter.

Single-pulse TMS was delivered with a figure-of-eight coil held at 45° from midline to induce an approximate posterolateral to anteromedial current. The hand motor hotspot and required TMS intensity were determined using standard procedures as outlined in **Supplementary** Fig. 1. To apply TUS and TMS concurrently, the ultrasound transducer was affixed to the center of the TMS coil using a custom-made 3D-printed clamp (**Fig. 1B**; Experiments I, II, & IV; Experiment III: see Fomenko et al., 2020). TMS was triggered 10 ms prior to the offset of TUS (**Fig. 1C**). Muscular activity was recorded in the first dorsal interosseous (FDI; Experiments I-III) or in the abductor pollicis brevis (APB; Experiment IV) via electromyography with surface adhesive electrodes using a belly-tendon montage (**Supplementary Table 1**).

In Experiments I, II, and IV, we used online neuronavigation with individual anatomical scans to support target selection and consistent TMS and TUS placement (Localite Biomedical Visualization Systems GmbH, Sankt Augustin, Germany; MRI specifications: **Supplementary Table 2**). Further, we recorded the position of TUS in Experiments I and II for post-hoc acoustic and thermal simulations.

### 2.3. Experiment I

On-target TUS was delivered to the left-hemispheric hand motor area to determine the effect of ultrasonic stimulation on corticospinal excitability. We introduced controls that improve upon the sole use of flip-over sham conditions used in prior work. First, we applied active control TUS to the right-hemispheric face motor area, allowing for the assessment of spatially specific effects while also better mimicking on-target peripheral confounds. In addition, we also included a sound-only sham condition that closely resembled the auditory confound (**Fig. 2**). Specifically, we generated a 1000 Hz square wave tone with 0.3 ms long pulses using MATLAB. We then added white noise at a signal-to-noise ratio of 14:1. This stimulus was administered to the participant via bone-conducting headphones (AfterShockz Trekz, TX, USA). Finally, we incorporated a baseline condition consisting solely of TMS.

**Fig. 2.**
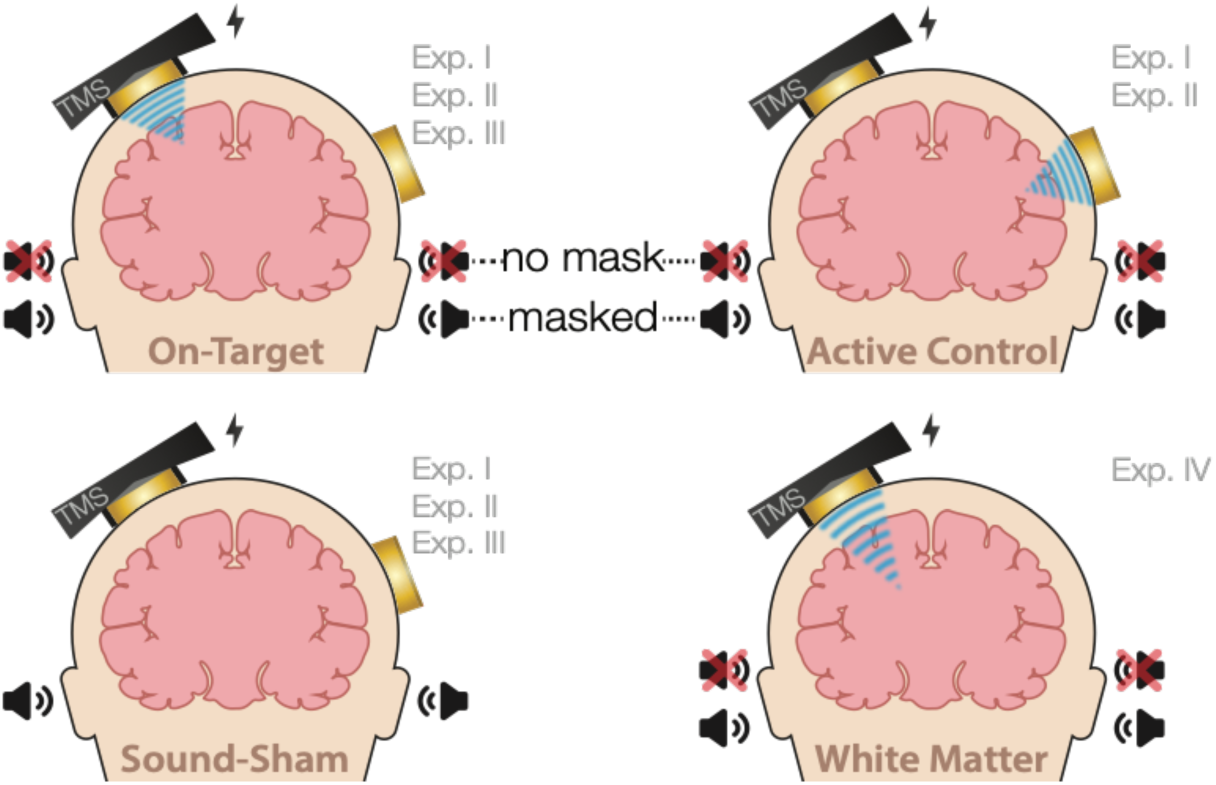
| Experimental conditions. On-target TUS of the left-hemispheric hand motor area (Exp. I-III), active control TUS of the right-hemispheric face motor area (Exp. I-II), sound-only sham (Exp. I-III), and inactive control TUS of the white matter ventromedial to the hand motor area (Exp. IV). Conditions involving TUS were presented both with and without auditory masking stimuli.

Ultrasonic stimulation was delivered at two pulse train durations (100/500 ms) and at two intensities (32.5/65 W/cm^2^ I_sppa_) to probe a potential dose-response effect. Additionally, with consideration of potentially audible differences between on-target and active control stimulation sites, we applied these conditions both with and without masking stimuli identical those used during sound-only sham. Auditory stimuli used for sound-sham and/or masking for each experiment are accessible here: https://doi.org/10.5281/zenodo.8374148. See **Supplementary** Fig. 3 for an overview of conditions and experimental timing for each experiment.

Conditions were administered in a single-blind inter-subject counterbalanced blocked design while participants were seated at rest. Ultrasound gel was used to couple both transducers to the participant’s scalp (Aquasonic 100, Parker Laboratories, NJ, USA). In total, participants completed 14 blocks of 20 trials each. Each trial lasted 6 ± 1 seconds. Two baseline measurements were completed, the first occurring as one of the first four blocks, and the second as one of the last four, to capture any general shift in excitability throughout the experiment. TMS was administered on every trial for a total of 280 single pulses.

### 2.4. Experiment II

To confirm and expand upon our findings from Experiment I we conducted a second, preregistered, experiment using the same main conditions and procedures, with a few adaptations (https://doi.org/10.17605/OSF.IO/HS8PT). The 2×2×2 design comprised of stimulation site (on-target/active control), stimulation intensity (6.35/19.06 W/cm^2^), and auditory masking (no mask/masked). We applied ultrasonic stimulation exclusively at an effective 500 ms pulse train duration. In this experiment, the same 1000 Hz square wave auditory stimulus was used for sound-only sham and auditory masking conditions. This stimulus was administered to the participant over in-ear headphones (ER-3C Insert Earphones, Etymotic Research, Illinois, USA). To better capture any baseline shift in excitability during the experiment, we presented conditions in a single-blind pseudorandomized order in which each consecutive set of 10 trials included each of 10 conditions once. Participants completed 25 trials per condition, resulting in 250 trials total.

To further probe a potential dose-response effect of stimulation intensity, we ran acoustic and thermal simulations (**Supplementary** Fig. 4). Here, we assessed the relationship between estimated intracranial intensities and perturbation of corticospinal excitability. While simulations were also run for Experiment I, its sample size was insufficient to test for intracranial dose-response effects.

Following the main experiment, we tested the efficacy of our masking stimuli with a forced-choice task wherein participants reported if they had received TUS for each condition, excluding baseline. Additionally, we investigated whether audible differences between stimulation sites were present during auditory masking (**Supplementary** Fig. 5).

### 2.5. Experiment III

We further characterized possible effects of auditory confounds on motor cortical excitability by administering varied auditory stimuli, both alongside on-target TUS and without TUS (i.e., sound-only sham). Auditory stimuli were either 500 or 700 ms in duration, the latter beginning 100 ms prior to TUS (**Supplementary** Fig. 3**.3**). Both durations were presented at two pitches. Using a signal generator (Agilent 33220A, Keysight Technologies), a 12 kHz sine wave tone was administered over speakers positioned to the left of the participant as in Fomenko and colleagues (2020). Additionally, a 1 kHz square wave tone with 0.5 ms long pulses was administered as in Experiments I, II, IV, and prior research (Braun et al., 2020) over noise-cancelling earbuds.

First, we investigated changes in corticospinal excitability from baseline following these auditory stimuli. Participants received 15 trials of baseline (i.e., TMS only) and 15 trials of each of the four sound-only sham stimuli. Conditions were presented in a blocked single-blind randomized order with participants seated at rest. An inter-trial interval of 5 seconds was used.

Next, we assessed whether applying on-target TUS during these auditory stimuli affected motor excitability. Here, TMS intensity was set to evoke a ∼1 mV MEP separately for each of the four sound-only sham conditions (**Supplementary** Fig. 1**.2**). To account for different applied TMS intensities between baseline and these conditions, we calculated *Relative MEP amplitude* by multiplying each trial by the ratio of applied TMS intensity to baseline TMS intensity. Participants received 15 trials of each auditory stimulus, once with on-target TUS and once as a sound-only sham. Ultrasound gel (Wavelength MP Blue, Sabel Med, Oldsmar, FL) and a 1.5 mm thick gel-pad (Aquaflex, Parker Laboratories, NJ, USA) were used to couple the transducer to the participants’ scalp. Conditions were presented in pairs of sound-sham and TUS for each auditory stimulus, counterbalanced between subjects. The order of the different auditory stimuli was randomized across participants.

### 2.6. Experiment IV

We further investigated the role of TUS audibility on motor excitability by administering stimulation to an inactive control site – the white matter ventromedial to the hand motor area. In doing so, TUS is applied over a homologous region of the scalp and skull without likely direct neuromodulation, thus allowing us to closely replicate the auditory confound while simultaneously isolating its effects.

Here, we probed whether the varying volume of the auditory confound at different stimulation intensities might itself impact motor cortical excitability. To this end, we applied stimulation at 4.34, 8.69, and 10.52 W/cm^2^ I_sppa_, or in effect, at three auditory confound volumes. We additionally applied stimulation both with and without a continuous auditory masking stimulus that sounded similar to the auditory confound. The stimulus consisted of a 1 kHz square wave with 0.3 ms long pulses. This stimulus was presented through wired bone-conducting headphones (LBYSK Wired Bone Conduction Headphones). The volume and signal-to-noise ratio of the masking stimulus were increased until the participant could no longer hear TUS, or until the volume became uncomfortable.

We administered conditions in a single-blind inter-subject randomized blocked design. Two blocks were measured per condition, each including 30 TUS-TMS trials and an additional 30 TMS-only trials to capture drifts in baseline excitability. These trials were applied in random order within each block with an inter-trial interval of 5 ± 1 seconds. Ultrasound gel (Aquasonic 100, Parker Laboratories, NJ, USA) and a ∼2-3 mm thick gel-pad were used to couple the transducer to the participant’s scalp (Aquaflex, Parker Laboratories, NJ, USA). During blocks with auditory masking, the mask was played continuously throughout the block. Following each block, participants were asked whether they could hear TUS (yes/no/uncertain).

### 2.7. Analysis

Raw data were exported to MATLAB, where MEP peak-to-peak amplitude was calculated using a custom script and confirmed by trial-level visual inspection. Trials where noise prevented an MEP to be sufficiently quantified were removed. Given the right-skewed nature of the raw MEP values, we performed a square root transformation to support parametric statistics. For visualization purposes, baseline corrected MEP amplitudes were also calculated.

Linear mixed-effects models (LMMs) were fitted using the lme4 package in R (Bates et al., 2015; R core team, 2021). Intercepts and condition differences (slopes) were allowed to vary across participants, including all possible random intercepts, slopes, and correlations in a maximal random effects structure (Barr et al., 2013). Statistical significance was set at two-tailed α = 0.05 and was computed with t-tests using the Satterthwaite approximation of degrees of freedom. For direct comparisons to a reference level (e.g., baseline), we report the intercept (*b*), standard error (*SE*), test-statistic (*t*), and significance (*p*). For main effects and interactions, we report the F statistic, significance, and partial eta squared. LMMs included square root transformed MEP peak-to-peak amplitude as the dependent variable, with the relevant experimental conditions and their interactions as predictors. Given the large number of baseline trials in Experiment IV (50% of total), the LMM testing effects of stimulation intensity and auditory masking instead included baseline corrected MEP amplitude as the dependent variable.

## 3. Results

### 3.1. Motor cortical inhibition is not specific to on-target TUS

We first corroborate previous reports of MEP suppression following 500 ms of TUS applied over the hand motor area (Experiments I-III; Fomenko et al., 2020; Legon, Bansal, et al., 2018; Xia et al., 2021). A LMM revealed significantly lower MEP amplitudes following on-target TUS as compared to baseline for Experiment I (*b* = –0.14, *SE* = 0.06, *t*(11) = –2.23, *p* = 0.047), Experiment II (*b* = –0.18, *SE* = 0.04, *t*(26) = –4.82, *p* = 6⋅10^-5^), and Experiment III (*b* = –0.22, *SE* = 0.07, *t*(15) = –3.08, *p* = 0.008).

However, corticospinal inhibition from baseline was also observed following control conditions. LMMs revealed significant attenuation of MEP amplitude following active control stimulation of the right-hemispheric face motor area (Experiment I: *b* = –0.12, *SE* = 0.05, *t*(11) = –2.29, *p* = 0.043; Experiment II: *b* = –0.22, *SE* = 0.04, *t*(26) = –5.60, *p* = 7⋅10^-6^), as well as after inactive control stimulation of the white matter ventromedial to the left-hemispheric hand motor area (Experiment IV: *b* = –0.14; *SE* = 0.04; *t*(11) = –3.09; *p* = 0.010). The same effect was observed following sound-only sham (Experiment I: *b* = –0.14; *SE* = 0.05; *t*(11) = –3.18; *p* = .009; Experiment II: *b* = –0.22; *SE* = 0.04; *t*(26) = –5.38; *p* = 1⋅10^-5^; Experiment III: 500ms-1kHz; *b* = –0.24; *SE* = 0.08; *t*(15) = –2.86; *p* = 0.012). These results suggest a spatially non-specific effect of TUS that is related to the auditory confound (**Fig. 3**).

**Fig. 3.**
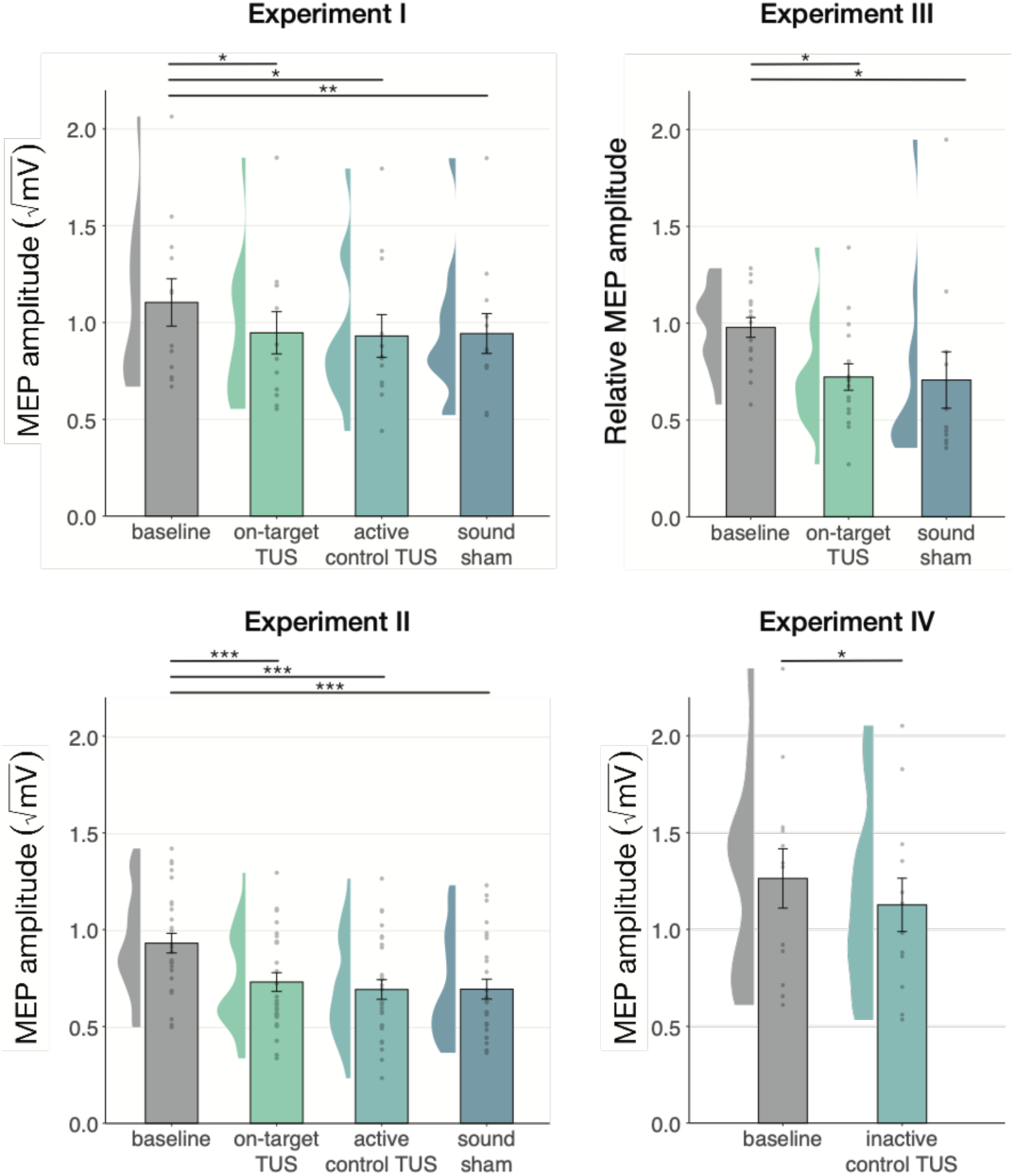
| Non-specific motor inhibitory effects of TUS. A significant suppression of MEP amplitude relative to baseline (gray) was observed for on-target TUS (green), but also for stimulation of a control region (cyan), and presentation of a sound alone (sound-sham; blue) indicating a spatially non-specific and sound-driven effect on motor cortical excitability. There were no significant differences between on-target and control conditions. Bar plots depict condition means, error bars represent standard errors, clouds indicate the distribution over participants, and points indicate individual participants. Square-root corrected MEP amplitudes are depicted for Experiments I, II, and IV, and *Relative MEP amplitude* is depicted for Experiment III (see ***Methods***). *p < 0.05, **p < 0.01, ***p < 0.001.

### 3.2. No dose-response effects of TUS on corticospinal inhibition

We further tested for direct ultrasonic neuromodulation by investigating a potential dose-response effect of TUS intensity (I_sppa_) on motor cortical excitability. First, we applied TUS at multiple free-water stimulation intensities (**Fig. 4C**). In Experiment I, a linear mixed model with the factor ‘intensity’ (32.5/65 W/cm^2^) did not reveal a significant effect of different on-target TUS intensities on motor excitability (*F*(1,11) = 0.47, *p* = 0.509, η_p_^2^ = 0.04). In Experiment II, a linear mixed model with the factors ‘stimulation site’ (on-target/active control), ‘masking’ (no mask/masked), and ‘intensity’ (6.35/19.06 W/cm^2^) similarly did not reveal an effect of stimulation intensity (*F*(1,50) = 1.29, *p* = 0.261, η_p_^2^ = 0.03). Importantly, there was no effect of stimulation site (*F*(1,168) = 1.75, *p* = 0.188, η_p_^2^ = 0.01), nor any significant interactions (all p-values > 0.1; all η_p_^2^ < 0.06). These results provide neither evidence for spatially specific neuromodulation when directly comparing stimulation sites, nor evidence for a dose-response relationship within the range of applied intensities.

**Fig. 4.**
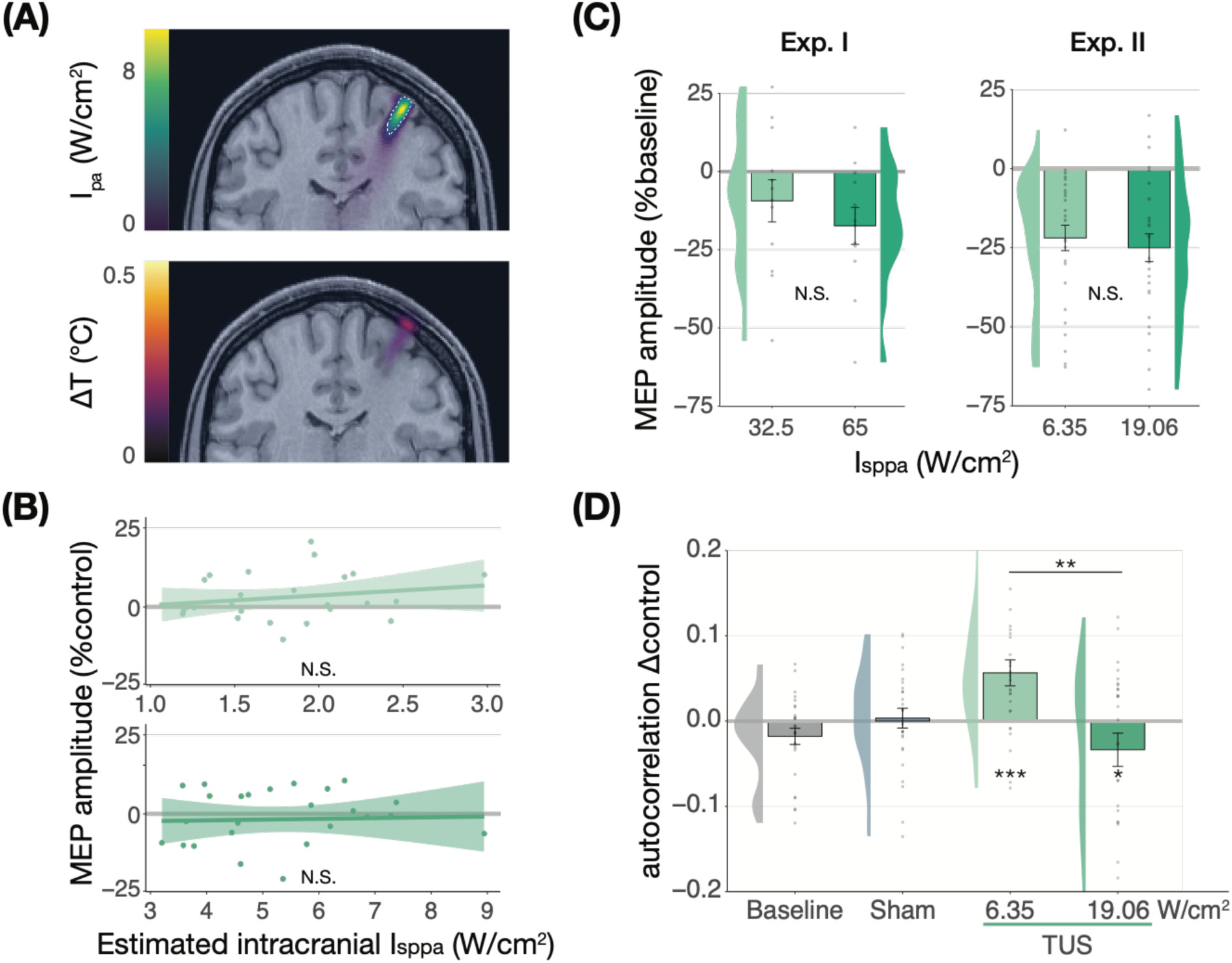
| No significant dose-response effects of TUS. **(A)** Acoustic (top) and thermal (bottom) simulations for a single subject in Experiment II. The acoustic simulation depicts estimated pulse-average intensity (I_pa_) above a 0.15 W/cm^2^ lower bound, with the dotted line indicating the full-width half-maximum of the pressure. The thermal simulation depicts maximum estimated temperature rise. **(B)** On-target TUS MEP amplitude as a percentage of active control MEP amplitude against simulated intracranial intensities at the two applied free-water intensities: 6.35 W/cm^2^ (top) and 19.06 W/cm^2^ (bottom). The shaded area represents the 95% CI, points represent individual participants. No significant intracranial dose-response relationship was observed. **(C)** There is no significant effect of free-water stimulation intensity on MEP amplitude. Values are expressed as a percentage of baseline MEP amplitude (square root corrected). Remaining conventions are as in Fig. 3. **(D)** Temporal autocorrelation, operationalized as the slope of the linear regression between trial *t* and its preceding baseline trial *t-*1, differed significantly as a function of stimulation site and intensity for masked trials. Individual points represent the differential autocorrelation compared to the active control site. Autocorrelation was not modulated during baseline or sound-only sham, but was significantly higher for on-target TUS at 6.35 W/cm^2^, and significantly lower for on-target TUS at 19.06 W/cm^2^ compared to active control TUS. *p < 0.05, **p < 0.01, ***p < 0.001.

However, it is likely that the effectiveness of TUS depends primarily on realized intracranial intensity, which we estimated with individualized 3D simulations (**Fig. 4A**). Yet, testing the relationship between estimated intracranial intensity and MEP amplitude change following on-target TUS similarly did not yield evidence for a dose-response effect (**Fig. 4B, Supplementary** Fig. 6).

Prior work has primarily focused on probing facilitatory or inhibitory effects on corticospinal excitability. Here, we also explored an alternative: how TUS might introduce noise to ongoing neural dynamics, rather than a directional modulation of excitability. Indeed, human TUS studies have often failed to show a global change in behavioral performance, instead finding TUS effects primarily around the perception threshold where noise might drive stochastic resonance (Butler et al., 2022; Legon et al., 2018). Whether the precise principles of stochastic resonance generalize from the perceptual domain to the current study is an open question, but it is known that neural noise can be introduced by brain stimulation (Van Der Groen & Wenderoth, 2016). It is likely that this noise is state-dependent and might not exceed the dynamic range of the intra-subject variability (Silvanto et al., 2007). Therefore, in an exploratory analysis, we exploited the natural structure in corticospinal excitability that exhibits as a strong temporal autocorrelation in MEP amplitude. Specifically, we tested how strongly the MEP on test trial *t* is predicted by the previous baseline trial *t-1*. As such, we quantified state-dependent autocorrelation between baseline MEP amplitude and MEP amplitude following on-target TUS, active control TUS, and sound-sham conditions (**Supplementary** Fig. 7). In brief, we found a significant interaction between previous baseline (*t-*1), stimulation site (on-target/active control), and intensity (6.35/19.06 W/cm^2^; *F*(1,30) = 12.10, *p* = 0.002, η_p_^2^ = 0.28) during masked trials. This interaction exhibited as increased autocorrelation for on-target TUS compared to active control TUS at 6.35 W/cm^2^ (i.e., lower TUS-induced noise; *F*(1,1287) = 13.43, *p* = 3⋅10^-4^, η_p_^2^ = 0.01), and reduced autocorrelation at 19.06 W/cm^2^ (i.e., higher noise; *F*(1,1282) = 5.76, *p* = 0.017, η_p_^2^ = 4⋅10^-3^; **Fig. 4D**). This effect was not only dependent upon intensity and stimulation site, but also dependent on the presence of auditory masking. As such, the effect was also observed in a four-way interaction of the previous baseline, site, intensity, and masking (**Supplementary** Fig. 7). These preliminary results might suggest that ultrasound stimulation can interact with ongoing neural dynamics by introducing temporally specific noise, rather than biasing the overall excitation/inhibition balance beyond its natural variation, but further work specifically designed to detect such effects is required.

### 3.3. Audible differences between stimulation sites do not underlie nonspecific inhibition

Stimulation over two separate sites could evoke distinct perceptual experiences arising from bone-conducted sound (Braun et al., 2020). To account for possible audibility differences between stimulation of on-target and active control sites in Experiments I and II, we also tested these conditions in the presence of a time-locked masking stimulus (**Fig. 5**). Following Experiment II, we additionally assessed the blinding efficacy of our masking stimuli, finding that the masking stimulus effectively reduced participant’s ability to determine whether TUS was administered to approximately chance level (**Supplementary** Fig. 5).

**Fig. 5.**
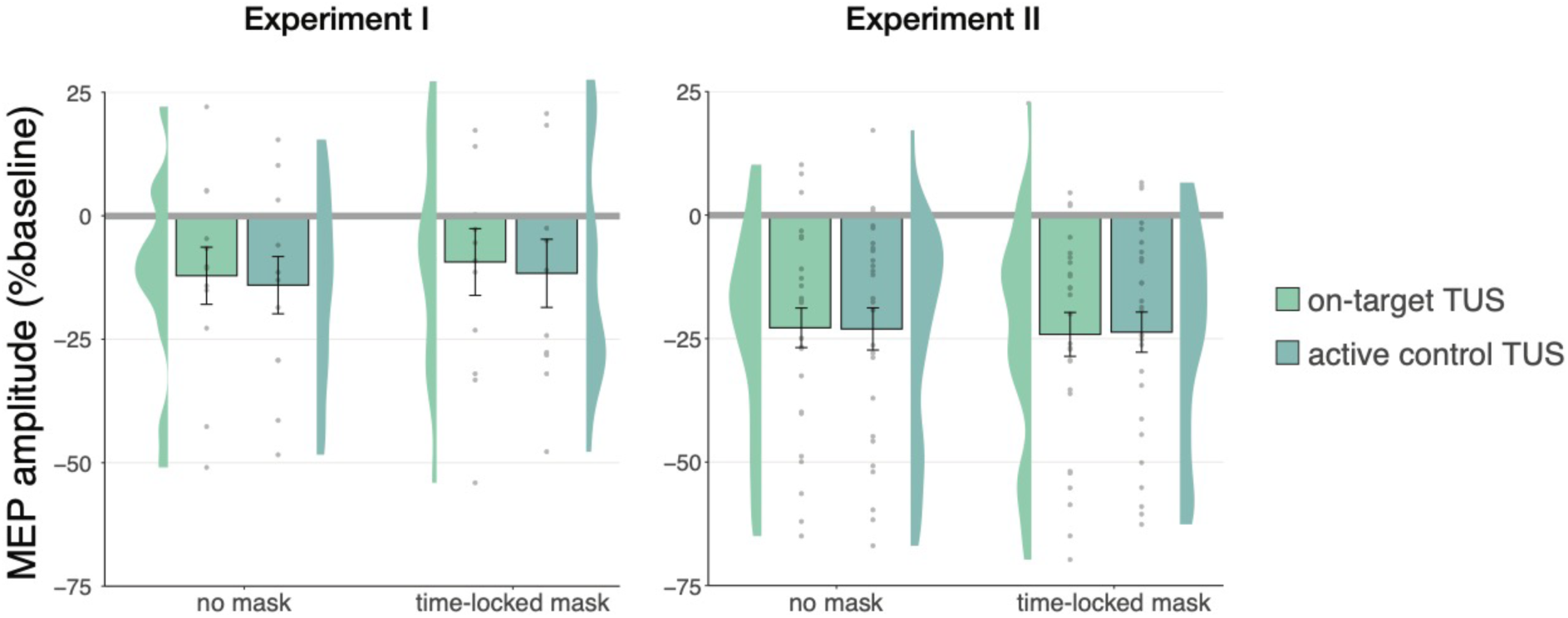
| No effects of time-locked masking. There were no significant effects of time-locked masking, indicating that audible differences between stimulation sites did not obscure or explain the absence of direct neuromodulation. Conventions are as in Figs. 3 and 4C.

In Experiment I, a linear mixed model with factors ‘masking’ (no mask/masked) and ‘stimulation site’ (on-target/active control) did not reveal a significant effect of masking (*F*(1,11) = 0.01, *p* = 0.920, η_p_^2^ = 1⋅10^-5^), stimulation site (*F*(1,11) = 0.15, *p* = 0.703, η_p_^2^ = 0.01), nor their interaction (*F*(1,11) = 1⋅10^-3^, *p* = 0.971, η_p_^2^ = 1⋅10^-4^). Similarly, in Experiment II, the linear mixed model described under the previous section revealed no significant main effect of masking (*F*(1,30) = 1.68, *p* = 0.205, η_p_^2^ = 0.05), nor any interactions (all p-values > 0.1; all η_p_^2^ < 0.06). These results indicate that an underlying specific neuromodulatory effect of TUS was not being obscured by audible differences between stimulation sites.

### 3.4. Sound-driven effects on corticospinal excitability

#### 3.4.1. Duration and pitch

Prior research has shown that longer durations of TUS significantly inhibited motor cortical excitability (i.e., ≥400 ms; Fomenko et al., 2020), while shorter durations did not. In Experiment I, we applied on-target, active control, and sound-sham conditions at shorter and longer durations to probe this effect. When directly comparing these conditions at different stimulus durations (100/500 ms), no evidence for an underlying neuromodulatory effect of TUS was observed, in line with our aforementioned findings. Instead, a linear mixed model with factors ‘condition’ (on-target/active control/sound-sham) and ‘stimulus duration’ (100/500 ms) revealed only a significant main effect of (auditory) stimulus duration, where longer stimulus durations resulted in stronger MEP attenuation (*F*(1,11) = 10.07, *p* = 0.009, η_p_^2^ = 0.48). There was no significant effect of condition (*F*(2,11) = 1.30, *p* = 0.311, η_p_^2^ = 0.19), nor an interaction between stimulus duration and condition (*F*(2,11) = 0.65, *p* = 0.543, η_p_^2^ = 0.11). These results further show that the auditory confound and its timing characteristics, rather than ultrasonic neuromodulation, underlies the observed inhibition of motor cortical excitability (**Fig. 6A**).

**Fig. 6.**
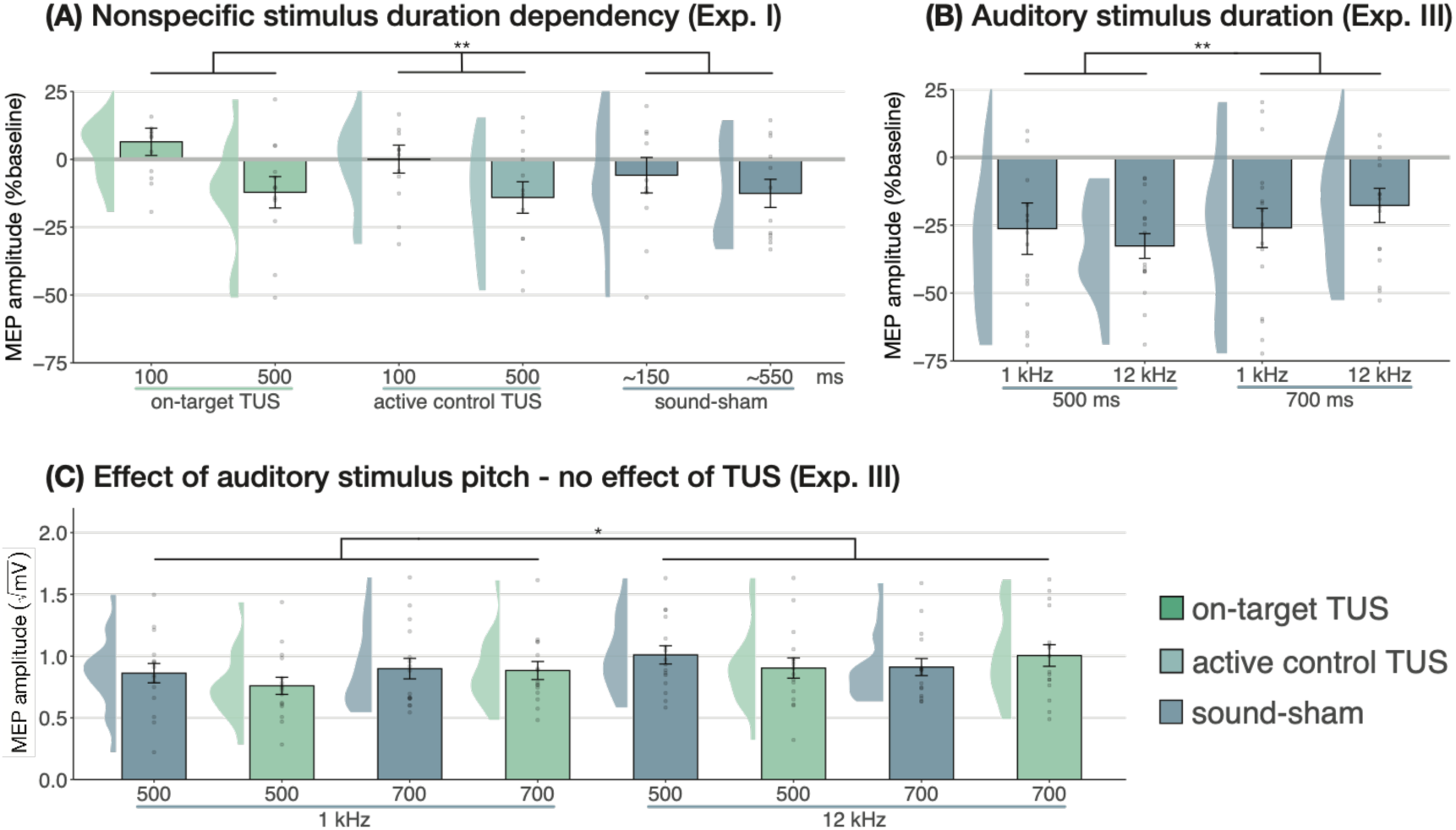
| Sound-driven effects on corticospinal excitability. **(A)** Longer (auditory) stimulus durations resulted in lower MEP amplitudes, regardless of TUS administration, indicating a sound-duration-dependency of motor inhibitory outcomes (Exp. I). **(B)** A significant effect of auditory stimulus duration was also observed in Experiment III. **(C)** The pitch of auditory stimuli also affected MEPs, where lower amplitudes were observed following a 1 kHz tone. There was no effect of TUS. Conventions are as in Figs. 3 and 4C.

We further tested auditory effects in Experiment III, where we administered sound-sham stimuli at four combinations of duration and pitch. A LMM with factors ‘duration’ (500/700 ms) and ‘pitch’ (1/12 kHz) revealed significantly lower MEPs following 500 ms auditory stimuli (**Fig. 6B**; duration: *F*(1,15) = 7.12, *p* = 0.017, η_p_^2^ = 0.32; pitch: *F*(1,15) = 0.02, *p* = 0.878, η_p_^2^ = 2⋅10^-3^; interaction: *F*(1,15) = 2.23, *p* = 0.156, η_p_^2^ = 0.13), supporting the role of auditory stimulus timing in perturbation of MEP amplitude.

Subsequently, ultrasonic stimulation was also administered alongside these four auditory stimuli. Here, a LMM with factors ‘auditory stimulus duration’ (500/700 ms), ‘pitch’ (1/12 kHz), and ‘ultrasonic stimulation’ (yes/no) revealed no significant effect of auditory stimulus duration in contrast to the first test (*F*(1,15) = 0.44, *p* = 0.517, η_p_^2^ = 0.03). However, a 1 kHz pitch resulted in significantly lower MEP amplitudes than a 12 kHz pitch (**Fig 6C**; *F*(1,15) = 4.94, *p* = 0.042, η_p_^2^ = 0.25). Importantly, we find no evidence for ultrasonic neuromodulation, where both on-target TUS and sound-sham reduced MEP amplitude from baseline (**Fig. 3**), and where applying on-target TUS did not significantly affect MEP amplitude as compared to sound-sham (*F*(1,15) = 0.42, *p* = 0.526, η_p_^2^ = 0.03; **Fig. 6C**). We observed a nonsignificant trend for the interaction between ‘ultrasonic stimulation’ and ‘auditory stimulus duration’ (*F*(1,15) = 4.22, *p* = 0.058, η_p_^2^ = 0.22). No trends were observed for the remaining interactions between these three factors (all η_p_^2^ < 0.06, *p* > 0.3). Taken together, these results do not provide evidence for direct ultrasonic neuromodulation but support the influence of auditory stimulation characteristics on motor cortical excitability.

#### 3.4.2. TUS audibility and confound volume

In Experiment IV, we applied TUS to an inactive target – the white matter ventromedial to the left-hemispheric hand motor area – both with and without a continuous auditory masking stimulus. MEP amplitudes did not significantly differ in baseline conditions regardless of whether a continuous sound was being played (*b* = 0.03, *SE* = 0.06, *t*(11) = 0.52, *p* = 0.616), indicating that continuous auditory stimulation alone might not be sufficient to inhibit MEP amplitude.

We additionally applied stimulation at multiple intensities to isolate the effect of auditory confound volume. A linear mixed model with factors ‘masking’ (no mask/masked) and ‘intensity’ (4.34/8.69/10.52 Wcm^-2^) with a random intercept and slope for each factor revealed a significant interaction (*F*(2,4038) = 3.43, *p* = 0.033, η_p_^2^ = 2⋅10^-3^) and an accompanying effect of ‘masking’ with lesser MEP attenuation when stimulation was masked (*F*(1,11) = 11.84, *p* = 0.005, η_p_^2^ = 0.52; **Fig. 7**). Follow-up comparisons revealed significantly less attenuation for masked stimulation at 4.34 W/cm^2^ intensity (*F*(1,11) = 13.02, *p* = 0.004, η_p_^2^ = 0.55), and a nonsignificant trend for the higher intensities (8.69 W/cm^2^: *F*(1,11) = 3.87, *p* = 0.077, η_p_^2^ = 0.27; 10.52 W/cm^2^: *F*(1,11) = 3.47, *p* = 0.089, η_p_^2^ = 0.24). In direct comparisons to baseline, all conditions resulted in a significant inhibition of MEP amplitude (all *t* < –3.36, all *p* < 0.007), with the exception of continuously masked stimulation at the lowest volume, with an intensity of 4.34 W/cm^2^ I_sppa_ (*b* = –0.06, *SE* = 0.03, *t*(11) = –2.04, *p* = 0.065).

**Fig. 7.**
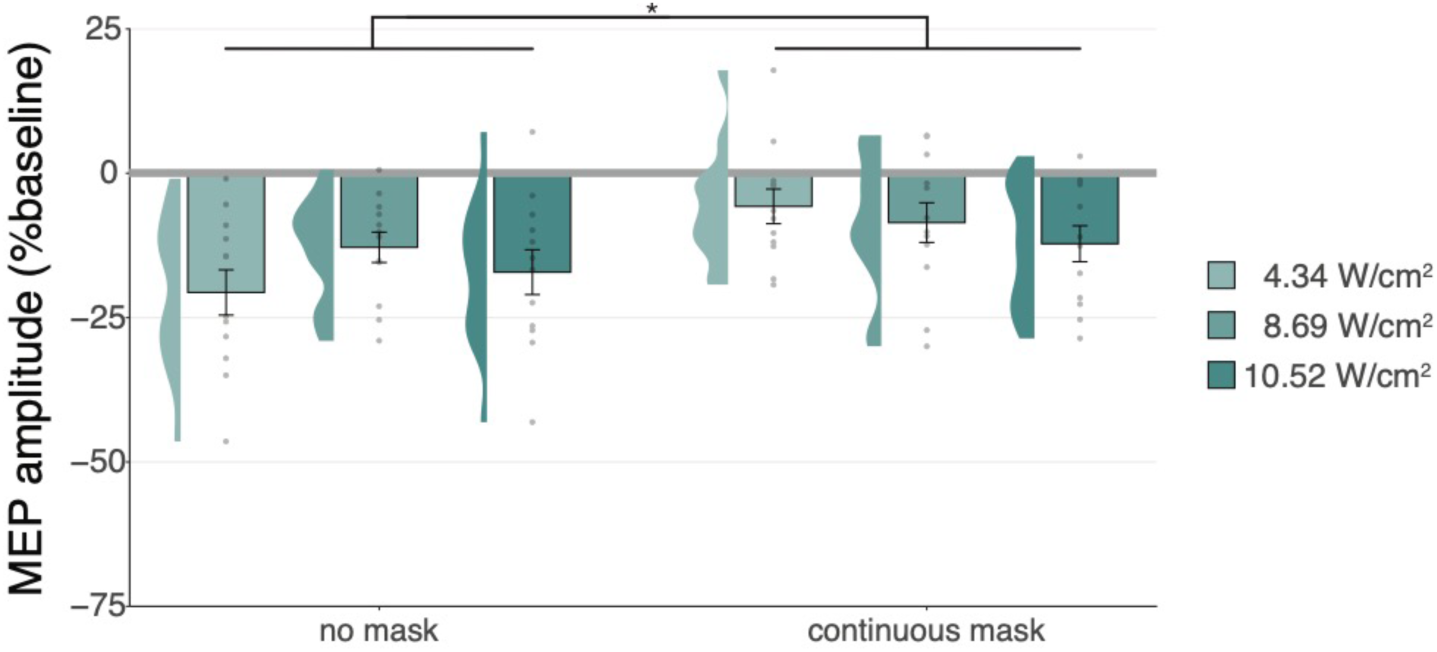
| Sound-driven effects on corticospinal excitability. Less MEP attenuation was measured during continuous masking, particularly for lower stimulation intensities (i.e., auditory confound volumes), pointing towards a role of TUS audibility in MEP attenuation.

The data indicate that continuous masking reduces motor inhibition, likely by minimizing the audibility of TUS, particularly when applied at a lower stimulation intensity (i.e., auditory confound volume). The remaining motor inhibition observed during masked trials likely owes to, albeit decreased, persistent audibility of TUS during masking. Indeed, MEP attenuation in the masked conditions descriptively scale with participant reports of audibility. This points towards a role of auditory confound volume in motor inhibition (***Supplementary*** Fig. 8). Nevertheless, one could instead argue that evidence for direct neuromodulation is seen here. This is unlikely for a number of reasons. First, white matter contains a lesser degree of mechanosensitive ion channel expression and there is evidence that neuromodulation of these tracts may occur primarily in the thermal domain (Guo et al., 2022; Sorum et al., 2021). Second, Experiment IV lacks sufficient inferential power in the absence of an additional control and must therefore be interpreted in tandem with Experiments I-III. These experiments revealed no evidence for direct neuromodulation using equivalent or higher stimulation intensities and directly targeting grey matter while also using multiple control conditions. Therefore, we propose that persistent motor inhibition during masked trials owes to continued, though reduced, audibility of the confound (***Supplementary*** Fig. 8). However, future work including an additional control (site) is required to definitively disentangle these alternatives.

#### 3.4.3. Preparatory cueing of TMS

We find that MEP attenuation results from auditory stimulation rather than direct neuromodulation. Two putative mechanisms through which sound cuing may drive motor inhibition have been proposed, positing either that explicit cueing of TMS timing results in compensatory processes that drive MEP reduction (Capozio et al., 2021; Tran et al., 2021), or suggesting the evocation of a startle response that leads to global inhibition (Fisher et al., 2004; Furubayashi et al., 2000; Ilic et al., 2011; Kohn et al., 2004; Wessel & Aron, 2013). Critically, we can dissociate between these theories by exploring the temporal dynamics of MEP attenuation. One would expect a startle response to habituate over time, where MEP attenuation would be reduced during startling initial trials, followed by a normalization throughout the course of the experiment. Alternatively, if temporally contingent sound-cueing of TMS drives inhibition, MEP amplitudes should decrease over time as the relative timing of TUS and TMS is being learned, followed by a stabilization at a decreased MEP amplitude once this relationship has been learned.

In Experiments I and II, linear mixed models with ‘trial number’ as a predictor show significant changes in MEP amplitude throughout the experiment, pointing to a learning effect. Specifically, in Experiment I, a significant reduction in MEP amplitude was observed across the first 10 trials where a 500 ms stimulus was delivered (*b* = –0.04, *SE* = 0.01, *t*(11) = –2.88, *p* = 0.015), following by a stabilization in subsequent blocks (*b* = –2⋅10^-4^, *SE* = 3⋅10^-4^, *t*(11) = –0.54, *p* = 0.601). This same pattern was observed in Experiment II, with a significant reduction across the first 20 trials (*b* = –0.01, *SE* = 3⋅10^-3^, *t*(26) = – 4.08, *p* = 4⋅10^-4^), followed by stabilization (*b* = 6⋅10^-5^, *SE* = 1⋅10^-4^, *t*(26) = 0.46, *p* = 0.650; **Fig. 8**). The data suggest that the relative timing of TUS and TMS is learned across initial trials, followed by a stabilization at a decreased MEP amplitude once this relationship has been learned. These results could reflect auditory cueing of TMS that leads to a compensatory expectation-based reduction of motor excitability.

**Fig. 8.**
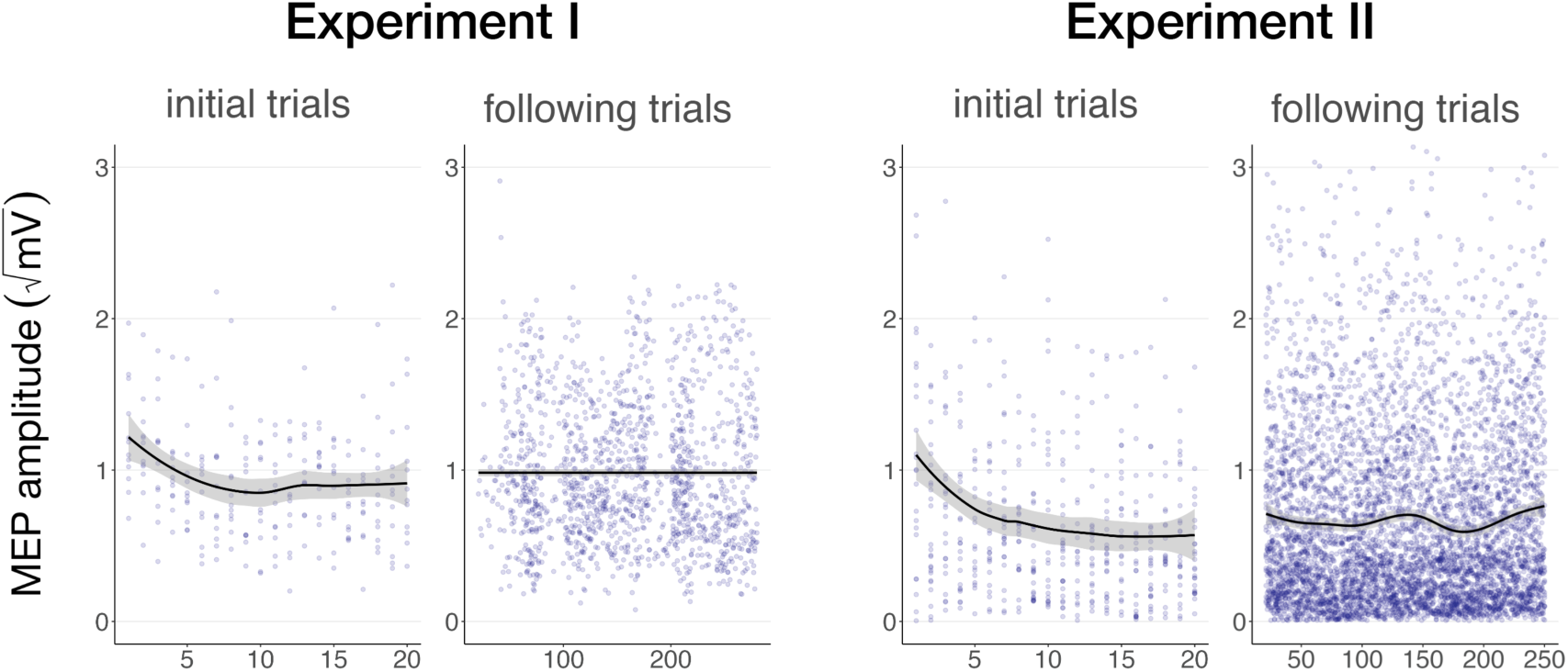
| Auditory cueing of TMS. There was a significant reduction in MEP amplitude when participants were first presented with a 500 ms stimulus (initial trials) in Experiment I (left) and Experiment II (right), following by a stabilization of MEP amplitude during the rest of the experiment (following trials), indicating a learning process by which TUS acts as a cue signaling the onset of TMS. The solid line depicts the loess regression fit, and the shaded area represents the 95% confidence interval.

## Discussion

In this study, we show the considerable impact of auditory confounds during audibly pulsed TUS in humans. We employed improved control conditions compared to prior work across four experiments, one preregistered, at three independent institutions. Here, we disentangle direct neuromodulatory and indirect auditory contributions during ultrasonic neuromodulation of corticospinal excitability. While we corroborated motor inhibitory effects of online TUS (Fomenko et al., 2020; Legon, Bansal, et al., 2018; Xia et al., 2021), we demonstrated that this inhibition also occurs with stimulation of a control region or presentation of a sound alone, suggesting that the auditory confound rather than direct ultrasonic neuromodulation drives inhibition. Further, no direct neuromodulatory effects on overall excitability were observed, regardless of stimulation timing, intensity, or masking. However, we note that an exploratory investigation of temporal dynamics indicated ultrasound might introduce noise to the neural system. Importantly, we found convincing evidence that characteristics of auditory stimuli do globally affect motor excitability, where auditory cueing of TMS pulse timing can affect measures of corticospinal excitability. This highlights the importance of explicit cueing in TMS experimental design. Most importantly, our results call for a reevaluation of earlier findings following audible TUS, and highlight the importance of suitable controls in experimental design (Bergmann & Hartwigsen, 2021; Siebner et al., 2022).

### No evidence for direct neuromodulation by TUS

Prior studies have highlighted sound-driven effects of TUS in behavioral and electrophysiological research (Airan & Butts Pauly, 2018; Braun et al., 2020; Guo et al., 2018; Johnstone et al., 2021; Sato et al., 2018). Here, we assessed whether the auditory confound of a conventional 1000 Hz pulsed protocol might underlie motor inhibitory effects, which are among the most robust and replicable human findings (Fomenko et al., 2020; Legon, Bansal, et al., 2018; Xia et al., 2021). While we successfully replicated this inhibitory effect, we found the same inhibition following stimulation of a motor control site (contralateral, active) and stimulation of a white-matter control site (ipsilateral, inactive; **Fig. 3**). This contrasts with a prior TUS-TMS study which found that TUS of the contralateral hand motor area did not change motor cortical excitability (Xia et al., 2021). Indeed, in all direct comparisons between on-target and control stimulation, no differences in excitability were observed, pointing towards a spatially nonspecific effect of TUS. Considering further inhibitory effects following administration of an auditory stimulus alone, the data suggest that online TUS motor inhibition is largely driven by the salient auditory confound, rather than spatially specific and direct neuromodulation. However, an exploratory analysis that tested for effects beyond a global shift in excitation-inhibition balance revealed that TUS might interact with ongoing neural dynamics by introducing dose-dependent noise (**Fig. 4D**).

We found no evidence of a dose-response relationship between TUS intensity (I_sppa_) and motor inhibition when applying stimulation at a wide range of intensities, nor when testing the relationship between simulated intracranial intensities and changes in excitability (**Fig. 4A-C**). Similarly, administration of a time-locked auditory masking stimulus that effectively reduced TUS detection rates did not provide evidence of direct effects being obscured by audible differences between conditions (**Fig. 5, Supplementary** Fig. 5). Taken together, this study presents no evidence for direct and spatially specific TUS inhibition of motor excitability when applying a clearly audible protocol, despite using improved control conditions, higher stimulation intensities, and a larger sample size than prior studies (Fomenko et al., 2020; Legon, Bansal, et al., 2018; Xia et al., 2021). Building on these results, the current challenge is to develop efficacious neuromodulatory protocols with minimal auditory interference. Efforts in this direction are already underway (Mohammadjavadi et al., 2019; Nakajima et al., 2022; Zeng et al., 2022).

### Sound-cued motor inhibition

Until now, it was unclear how TUS induced motor inhibition in humans. Here, we show that this inhibition is caused by peripheral auditory stimulation. It is well-known that MEPs are sensitive to both sensory and psychological factors (Duecker et al., 2013). For example, several studies find MEP attenuation following a startling auditory stimulus (Fisher et al., 2004; Furubayashi et al., 2000; Ilic et al., 2011; Kohn et al., 2004; Wessel & Aron, 2013), and have demonstrated the impact of stimulus duration and volume on this inhibition (Furubayashi et al., 2000). It is possible that a similar mechanism is at play for audible TUS protocols. Indeed, we observed modulation of motor cortical excitability dependent upon the characteristics of auditory stimuli, including their duration and timing (**Fig. 6A-B**), their pitch/frequency (**Fig. 6C**), and whether the confound was audible in general, including perceived volume (**Fig. 7, Supplementary** Fig. 8).

One possible interpretation of the observed MEP attenuation is that the auditory confound acts as a salient cue to predict the upcoming TMS pulse. Prediction-based attenuation has been reported in both sensory and motor domains (Ford et al., 2007; Tran et al., 2021). For example, MEPs are suppressed when the timing of a TMS pulse can be predicted by a warning cue (Capozio et al., 2021; Tran et al., 2021). In the current experimental setup, participants could also learn the relative timing of the auditory stimulus and the TMS pulse. Indeed, we observe MEP attenuation emerge across initial trials as participants learn when to expect TMS, until a stable (i.e., learned) state is reached (**Fig. 8**). Moreover, no motor inhibition was observed when TUS onset was inaudible or when stimulation timing was potentially too fast to function as a predictive cue (100 ms). Taken together, a parsimonious explanation is expectation-based inhibition of TMS-induced MEPs. This inhibitory response might either reflect inhibition of competing motor programs – a component of motor preparation – or a homeostatic process anticipating the TMS-induced excitation (Capozio et al., 2021; Tran et al., 2021).

### Limitations

The precise biomolecular and neurophysiological mechanisms underlying ultrasonic neuromodulation remain under steadily progressing investigation (Weinreb & Moses, 2022; Yoo et al., 2022). A shared interpretation is that mechano-electrophysiological energy transfer is proportional to acoustic radiation force, and thus proportional to stimulation intensity. Accordingly, one could argue that the TUS dose in the present study could have been insufficient to evoke direct neuromodulation. Indeed, despite the applied intensities exceeding prior relevant human work (Fomenko et al., 2020; Legon, Bansal, et al., 2018; Xia et al., 2021) the total applied neuromodulatory doses are relatively low as compared to, for example, repetitive TUS protocols (rTUS) in animal work (Folloni et al., 2019; Verhagen et al., 2019) or recent human studies (Nakajima et al., 2022).

Alternatively, insufficient neural recruitment could be attributed to stimulation parameters other than intensity. If so, the absence of direct neuromodulation across these experiments might not generalize to parameters outside the tested set. For example, while we replicated and extended prior work targeting the hand motor area at ∼30 mm from the scalp (Fomenko et al., 2020; Legon, Bansal, et al., 2018; Xia et al., 2021), other studies have suggested that the optimal stimulation depth to engage the hand motor area may be more superficial (Osada et al., 2022; Siebner et al., 2022).

One might further argue that the TMS hotspot provides insufficient anatomical precision to appropriately target the underlying hand muscle representation with TUS. The motor hotspot may not precisely overly the cortical representation of the assessed muscle due to the increased coil-cortex distance introduced by the TUS transducer. This distance, and the larger TMS coils required to evoke consistent MEPs, results in a broad electric field that is substantially larger than the TUS beam width (e.g., 6 mm for 250 kHz; Fomenko et al., 2020; Legon, Bansal, et al., 2018). Thus, it is possible that a transducer aligned with the center of the TMS coil may not be adequate. Nevertheless, we note that previous work utilizing a similar targeting approach has effectively induced changes in corticospinal motor excitability (Zeng et al., 2022). We also note that our stimulation depth and targeting procedures were comparable to all prior TUS-TMS studies, and that our simulations confirmed targeting (**Fig. 3A, Supplementary** Fig. 4). In summary, our main finding that the auditory confound drove motor inhibition in the present study, and likely had an impact in previous studies, holds true.

### Considerations and future directions

Crucially, our results do not provide evidence that TUS is globally ineffective at inducing neuromodulation. While the present study and prior research highlight the confounding role of indirect auditory stimulation during pulsed TUS, there remains strong evidence for the efficacy of ultrasonic stimulation in animal work when auditory confounds are accounted for (Mohammadjavadi et al., 2019), or in controlled in-vitro systems such as an isolated retina, brain slices, or neuronal cultures in which the auditory confound carries no influence (Menz et al., 2013; Tyler et al., 2018).

It follows that where an auditory confound could be expected, appropriate control conditions are critical. These controls could involve stimulating a control region, and/or including a matched sound-only sham. In parallel, or perhaps alternatively, the impact of this confound can be mitigated in several ways. First, we recommend that the influence of auditory components be considered in transducer design and selection. Second, masking the auditory confound can help blind participants to experimental conditions. Titrating auditory mask quality per participant to account for intra– and inter-individual differences in subjective perception of the auditory confound would be beneficial. Here, the method chosen for mask delivery must be considered. While bone-conducting headphones align with the bone conduction mechanism of the auditory confound, they might not deliver sound as clearly as in-ear headphones or speakers. Nevertheless, the latter two rely on air-conducted sound. Notably, in-ear headphones could even amplify the perceived volume of the confound by obstructing the ear canal. Importantly, even when using masking stimuli, auditory stimulation could still influence cognitive task performance, among other measures. Alternative approaches could circumvent auditory confounds by testing deaf subjects, or perhaps more practically by ramping the ultrasonic pulse to minimize or even eliminate the auditory confound. This approach still requires validation and will only be relevant for protocols with pulses of sufficient duration. Here, one can expect that the experimental control required to account for auditory confounds might also hold for alternative peripheral effects, such as somatosensory confounds. Longer pulse durations are common in offline rTUS paradigms (Zeng et al., 2022), with more opportunity for inaudible pulse shaping and the added benefit of separating the time of stimulation from that of measurement. However, appropriate control conditions remain central to make inferences on interventional specificity.

## Conclusion

Transcranial ultrasonic stimulation is rapidly gaining traction as a promising neuromodulatory technique in humans. For TUS to reach its full potential we must identify robust and effective stimulation protocols. Here, we demonstrate that one of the most reliable findings in the human literature – online motor cortical inhibition during a 1000 Hz pulsed protocol – primarily stems from an auditory confound rather than direct neuromodulation. Instead of driving overall inhibition, we found preliminary evidence that TUS might introduce noise to ongoing neural dynamics. Future research must carefully account for peripheral confounding factors to isolate direct neuromodulatory effects of TUS, thereby enabling the swift and successful implementation of this technology in both research and clinical settings.

## Data availability

Data and code for the current study can be accessed by reviewers <u>here</u>. An explanation of data access during the review processes can be found on <u>this page</u>.

**Fig. 6.**
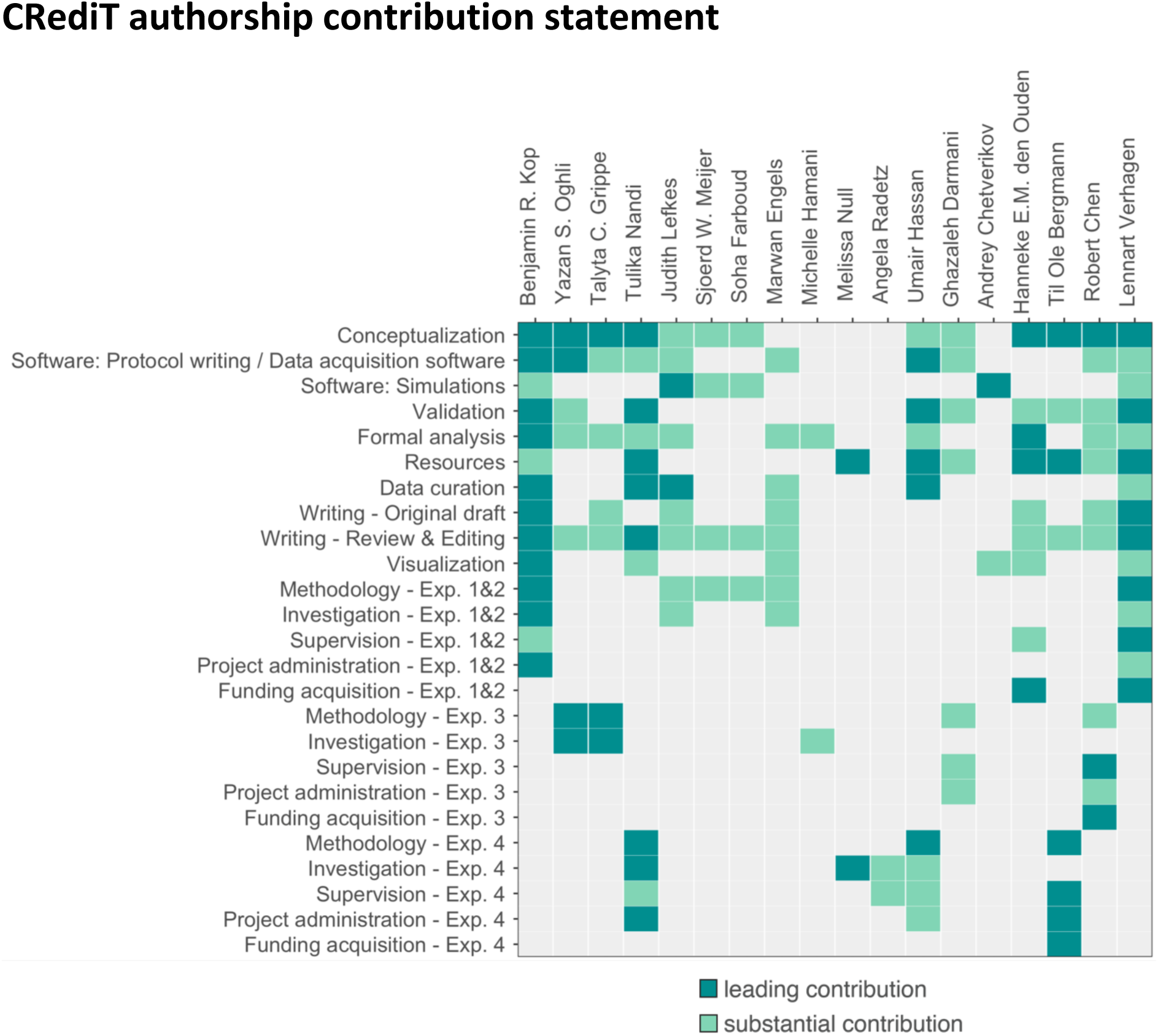
| Contribution diagram. This figure depicts the involvement of each author using the CRediT taxonomy (Brand et al., 2015) and categorizes their contributions according to three levels represented by color: ‘none’ (gray), ‘substantial contribution’ (light green), ‘leading contribution’ (dark green).

## Acknowledgements

Experiments I & II were supported by the Dutch Research Council (NWO), awarding VIDI fellowships to L.V. (18919) and H.E.M.d.O. (452-17-016), and the Topsector programme Holland High Tech (HiTMaT-38H3). We thank Brittany van Beek for assistance with data acquisition and administration, Sarmad Peymaei for contributions to technical piloting, Norbert Hermesdorf for developing the 3D-printed TUS-TMS clamp, and Gerard van Oijen and Pascal de Water for their technical support. Experiment III was funded by the Canadian Institutes of Health Research (FDN 154292, ENG 173742) and the Natural Science and Engineering Research Council of Canada (RGPIN-2020-04176, RTI-2020-0024). Experiment IV and T.N. were supported by a grant from the Boehringer Ingelheim Foundation to T.O.B.

## Author ORCIDs

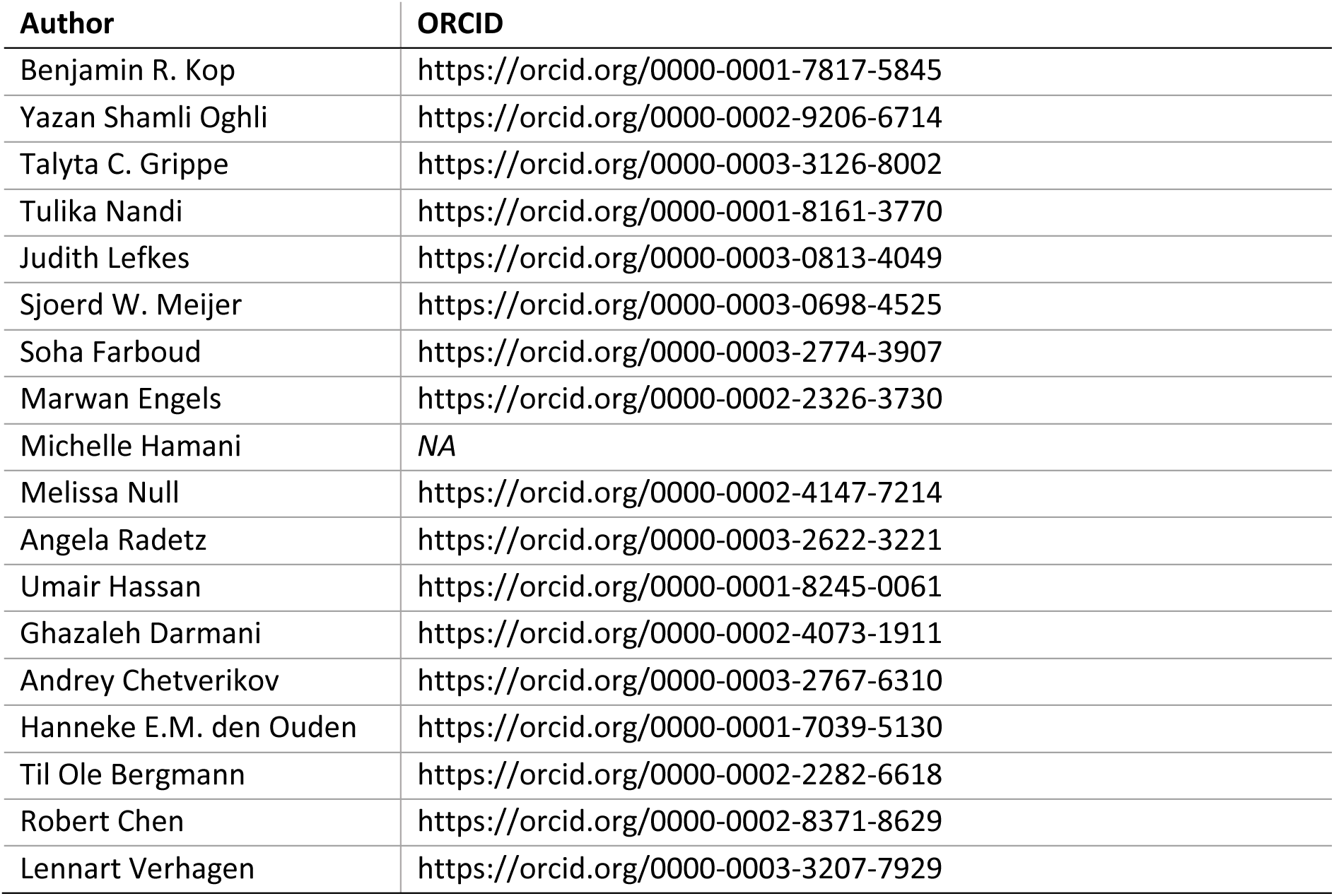

## Appendix A. Supplementary information

Supplementary tables and figures can be found in Appendix A.

