## Appendix A - Supplementary Material for "Auditory confounds can drive online effects of transcranial ultrasonic stimulation in humans"

For questions, please contact the corresponding author.

### Supplementary Tables

#### Supplementary Table 1 | Transducer specifications

| Exp. | Model number | *N elements* | Radius of curvature | Aperature diameter | Width | Solid water coupling |
|---|---|---|---|---|---|---|
| I | CTX500-006 | 2 | 63.2 | 45.0 | ~16 | yes |
| II | CTX250-014 | 2 | 63.2 | 45.0 | ~16 | yes |
| III | H246-01 | 2 | 0 (flat) | 33.6 | 10 | no |
| IV | CTX250-005 | 2 | 63.2 | 45.0 | 16.45 | yes |

This table describes transducer specifications used with a Sonic Concepts TPO (Bothell, WA, USA).

#### Supplementary Table 2 | Electromyography

| Experiment | Muscle | Amplification | Filtering | Digital sampling rate | Software |
|---|---|---|---|---|---|
| I & II | FDI | 1000 gain[1] | 1 – 1000 Hz[1] | 5 kHz[2] | Signal version 7.05[3] |
| III | FDI | 1000 gain[4] | 20 – 2500 Hz[4] | 5 kHz[2] | Signal version 6.04[3] |
| IV | APB | 10 gain[5] | <1250 Hz[5] | 5 kHz[5] | BEST Toolbox[6] |

[1]D440-2, Digitimer Ltd., Hertfordshire, UK; [2]Micro 1401, CED, Cambridge, UK; [3]CED, Cambridge, UK; [4]Intronix Technologies Corporation [model: 2024F], Bolton, Canada; [5]Bittium NeurOne Tesla System, Bittium Biosignals Ltd., Finland; [6]Hassan et al., 2022.

This table describes the EMG acquisition parameters used to measure muscular activity in the first dorsal interosseous (FDI) or abductor pollicis brevis (APB) using a belly-tendon montage.

#### Supplementary Table 3 | MRI acquisition parameters

| Experiment | Scan | TR | TE | FoV read | FoV phase | voxel | N slices |
|---|---|---|---|---|---|---|---|
| I & II | T1w[1] | 2700 ms | 3.69 ms | 230 mm | 128.1% | 0.9 mm iso | 224 |
| | T2w[1] | 3200 ms | 408 ms | 230 mm | 128.1% | 0.9 mm iso | 224 |
| IV | T1w[2] | 2700 ms | 3.7 ms | 260 mm | 88.9% | 0.9 mm iso | 224 |

[1]3T Siemens Skyra MRI scanner (Siemens Medical Solutions, Erlangen, Germany) with a 32-channel head coil. [2]3T Siemens Prisma MRI scanner with a 32-channel head coil.

This table shows the MRI acquisition parameters used in Experiments I, II, and IV. Anatomical T1w scans were used for online neuronavigation. For Experiments I and II, both T1w and T2w scans were used for post-hoc acoustic and thermal simulations.

#### Supplementary Table 4 | Properties for acoustic and thermal simulations (Exp. I-II).

| | Density (kg/m$^3$) | Sound speed (ms) | Attenuation coefficient | Thermal conductivity (W/m/°C) | Specific heat capacity (J/kg/°C) |
|---|---|---|---|---|---|
| Water | 994 | 1500 | 0.0 | 0.60 | 4178 |
| Scalp | 1090 | 1610 | 0.4 | 0.37 | 3391 |
| Skull | 1850 | 2800 | 8.0 | 0.32 | 1313 |
| Brain | 1046 | 1546 | 0.6 | 0.51 | 3630 |

This table describes the acoustic and thermal properties assigned to each component included in simulations. See **Supplementary Figure 4**.

### Supplementary Figures

**Supplementary Figure 1.1 |** TMS hotspot and intensity determination.

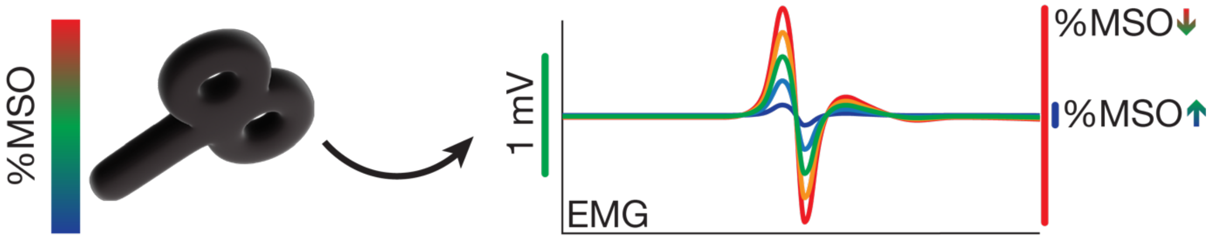

Percentage of maximum stimulator output (%MSO) is determined by finding the motor hotspot and then adjusting %MSO until a stable 1 mV MEP is observed (Experiments I-III) or until a MEP of minimally 0.05 mV is observed on at least half of trials (Experiment IV).

##### *Experiments I & II*

Single-pulse TMS was delivered with a 97 mm figure-of-eight MC-B70 coil powered by a MagPro X100 + MagOption stimulator (MagVenture, Farum, Denmark) using a biphasic pulse shape. The motor hotspot for the first dorsal interosseous (FDI) was determined by positioning the TMS coil over the hand motor area as identified with neuronavigation software, and iteratively adjusting TMS intensity until a consistent MEP was observed of approximately 1 mV amplitude. After probing locations in a ~2 cm radius, the location resulting in the highest and most stable amplitude MEPs was set as the final stimulation site. Next, the percentage of maximum stimulator output (%MSO) was adjusted until an average MEP amplitude of ~1 mV was observed across approximately ten trials. This intensity was used throughout the rest of the experiment. Both TMS and TUS were externally triggered using Signal version 7.05 (CED, Cambridge, UK). The mean percentage of administered TMS maximum stimulator output was 80.4 ± 12.5% for Experiment I, and 88.0 ± 9.3% for Experiment II. These high stimulation intensities are to be expected, given the offset of TMS from the scalp owing to the ultrasound transducer and the TMS current dissipating with distance (O'Shea & Walsh, 2007).

##### *Experiment III*

Single-pulse TMS was delivered via a 70 mm figure-of-eight coil powered by a Magstim $200^2$ stimulator (Magstim, Whitland, Dyfed, UK). The coil was held at 45° from midline to induce an approximate posterolateral to anteromedial using a monophasic pulse shape. Optimal positioning was determined by placing the TMS coil over the left-hemispheric hand motor area and moving the coil in increments of 0.5 cm until a clear MEP was observed over the FDI. Following assessment of optimal coil flatness and orientation, the location and orientation of TMS were marked on the scalp to support consistent placement throughout the experiment. At this site, the minimal intensity required to evoke an average MEP of ~1 mV across ten trials was determined. The mean %MSO was 73.7 ± 11.1% for the baseline condition (i.e., TMS only). The mean %MSO require to evoke a ~ 1mV MEP during the four sound-sham conditions were significantly higher at 77.4 ± 9.4% [1 kHz, 500 ms, $t(15) = -4.64$, $p = 3 \cdot 10^{-4}$], 77.6 ± 10.1% [1 kHz, 700 ms, $t(15) = -4.84$, $p = 2 \cdot 10^{-4}$], 76.9 ± 10.6% [12 kHz, 500 ms, $t(15) = -3.76$, $p = 0.002$], and 77.2 ± 9.8% [12 kHz, 700 ms, $t(15) = -4.09$, $p = 9.7 \cdot 10^{-4}$, see Supplementary Fig. 1.2]. To administer stimulation at high temporal precision, both TUS and TMS were externally triggered using Signal 6.04 (CED, Cambridge, UK).

*Experiment IV*

The TMS stimulator, coil, pulse shape, and orientation were identical to Experiments I & II, excluding the final four participants in which the Cool-B35 HO coil (Magventure, Farum, Denmark) was used during preparatory measures to allow more precise estimation of the motor hotspot, while always the same TMS coil (MC-B70) was used during the combined TUS-TMS experiment.

The motor hotspot for the abductor pollicis brevis (APB) was determined by positioning the TMS coil over the hand motor area as identified with neuronavigation software, and systematically searching for the largest and most consistent MEP. An automated adaptive staircase procedure with an amplitude threshold of 0.05 mV was used to determine the resting motor threshold (rMT). We subsequently estimated the dose response curve by applying 20 pulses each at 80, 90, 100, 110, 120, and 130% rMT. In most cases we were unable to reach the plateau of the dose-response curve due to the high motor thresholds resulting from the offset caused by the ultrasound transducer. We aimed to use 120% rMT (n =3). However, if this intensity surpassed 100% MSO, we opted for 100% MSO instead (n = 9). The mean %MSO was 94.5 ± 10.5%. The TMS intensities required in this experiment were higher than those required in Experiment I-II using the same TMS coil, though still within approximately one standard deviation. This is likely due to the use of a gel pad, which introduces more distance between the TMS coil and the scalp, thus requiring a higher TMS intensity to evoke the same motor activity. To administer stimulation at high temporal precision, both TUS and TMS were externally triggered via a bossdevice (sync2brain, Tübingen, Germany) using the BEST Toolbox (Hassan et al., 2022).

**Supplementary Figure 1.2 |** TMS intensities in Experiment III

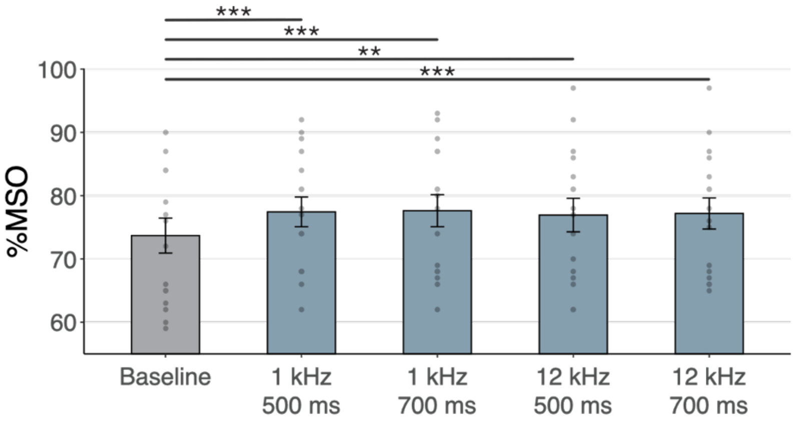

Percentage maximum stimulator output (%MSO) required in Experiment III to evoke a 1 mV MEP at baseline and during the four sound-sham conditions. Each sound-sham conditions required a significantly higher %MSO than baseline to evoke a 1 mV MEP. *p < 0.05, **p < 0.01, **p < 0.001.

**Supplementary Figure 2 | Simulated intracranial indices (Exp. I-II).**

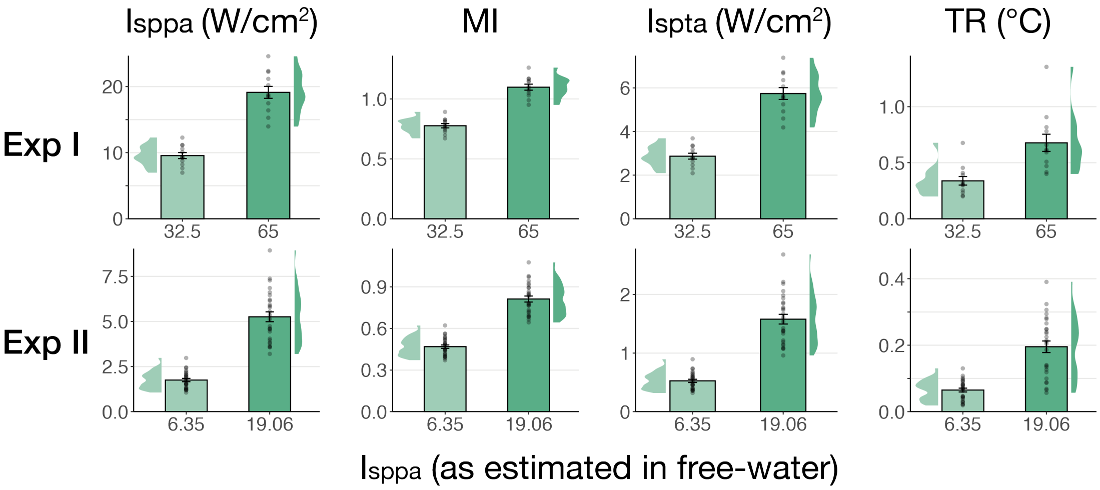

Simulated indices for Experiment I (top) and Experiment II (bottom) for both applied free-water stimulation intensities including spatial-peak pulse- and temporal-average intensity ($I_{sppa}$, $I_{spta}$), the mechanical index (MI), and peak thermal rise (TR).

**Supplementary Figure 3.1 |** Overview of experimental conditions

| | stimulation condition | free-water intensity ($I_{sppa}$) | auditory masking | TUS duration |
|---|---|---|---|---|
| **Exp. I** | on-target TUS<br>active control TUS<br>sound-sham<br>baseline | 32.5 W/cm$^2$<br>65 W/cm$^2$ | time-locked to TUS<br>no mask<br>masked | 100 ms<br>500 ms |
| **Exp. II** *preregistered* | on-target TUS<br>active control TUS<br>sound-sham<br>baseline | 6.35 W/cm$^2$<br>19.06 W/cm$^2$ | time-locked to TUS<br>no mask<br>masked | 500 ms |
| **Exp. III** | on-target TUS<br>sound-sham<br>baseline | 9.26 W/cm$^2$ | time-locked to TUS<br>masked<br>1 kHz, 500 ms<br>1 kHz, 700 ms<br>12 kHz, 500 ms<br>12 kHz, 700 ms | 500 ms |
| **Exp. IV** | inactive control TUS<br>baseline | 4.34 W/cm$^2$<br>8.69 W/cm$^2$<br>10.52 W/cm$^2$ | continuous masking<br>no mask<br>masked | 400 ms |

For each experiment (row), core experimental conditions are noted. In Experiments I and II, main conditions included on-target TUS (left-hemispheric hand motor area), active control TUS (right-hemispheric face motor area), sound-sham (auditory stimulation alone), and baseline (TMS only). Here, TUS was applied at two stimulation intensities, and both with and without a time-locked auditory masking stimulus. We report the spatial-peak pulse-averaged intensity in free-water without attenuation from biological tissue. In Experiment I, TUS was further administered at both 100 and 500 ms stimulus durations. Experiment II is a preregistered study to confirm and extend the findings of Experiment I. In Experiment III, four auditory stimuli of varying pitch and duration were administered both in isolation (sound-sham) and alongside on-target TUS. In Experiment IV, inactive control TUS was administered to the white matter ventromedial to the hand motor area at three intensities (i.e., auditory confound volumes) both with and without a continuous auditory masking stimulus.

**Supplementary Figure 3.2 |** Conditions and timing of Experiments I and II

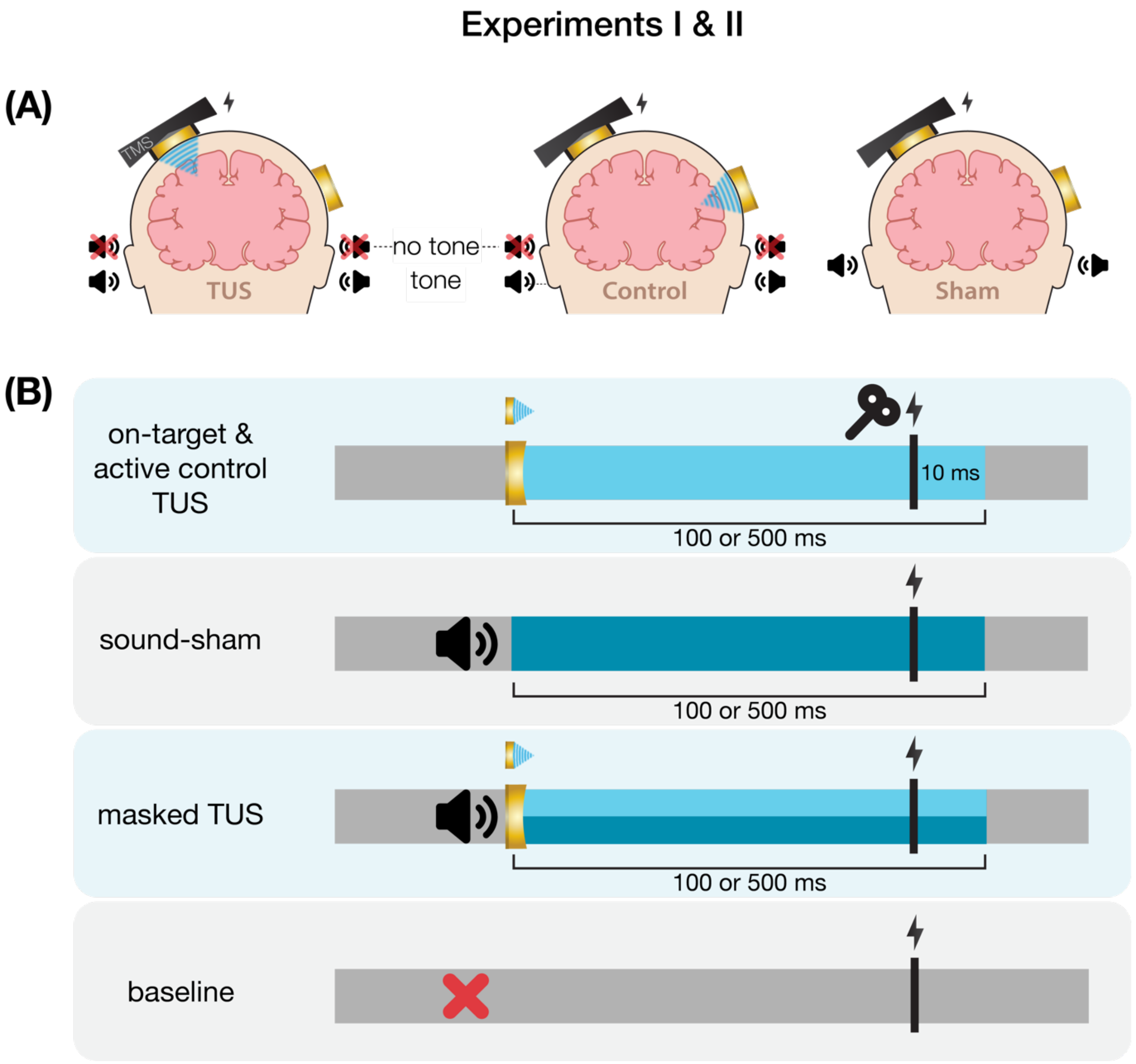

**(A)** On-target left-hemispheric hand area stimulation (left) and active control right-hemispheric face area stimulation (middle) were applied both with and without a masking tone. The sham condition (right) consisted solely of an identical tone. **(B)** On-target and active control TUS were administered for 100 ms [Experiment I] and 500 ms [Experiments I & II] with TMS applied 10 ms prior to TUS-offset. In Experiment I, the auditory stimulus administered during sound-sham and masked conditions began ~50 earlier and was 100 ms longer than TUS, while in Experiment II the auditory stimulus was precisely timed to match TUS. Baseline measurement involved no intervention other than TMS.

**Supplementary Figure 3.3 |** Conditions and timing of Experiment III

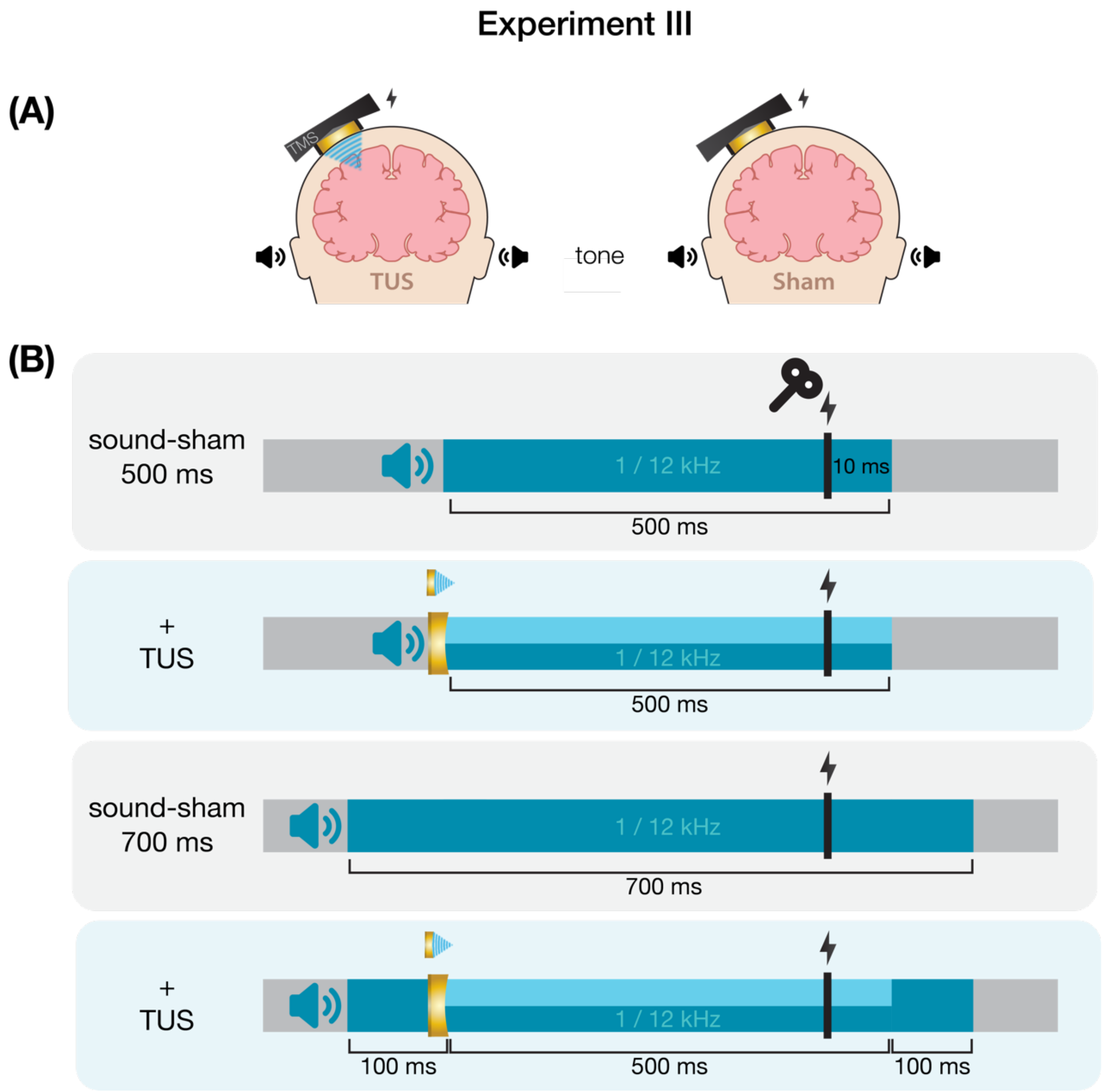

**(A)** On-target TUS consisted of left-hemispheric hand motor area stimulation in concurrence with an auditory stimulus (left), while sound-sham consisted solely of an auditory stimulus (right). **(B)** Auditory stimuli were delivered at both 1 and 12 kHz for either 500 or 700 ms. TUS was delivered for 500 ms. TMS was delivered 10 ms prior to TUS offset (i.e., following 490 ms of TUS), and at the same timing in the absence of TUS.

**Supplementary Figure 3.4 |** Conditions and timing of Experiment IV

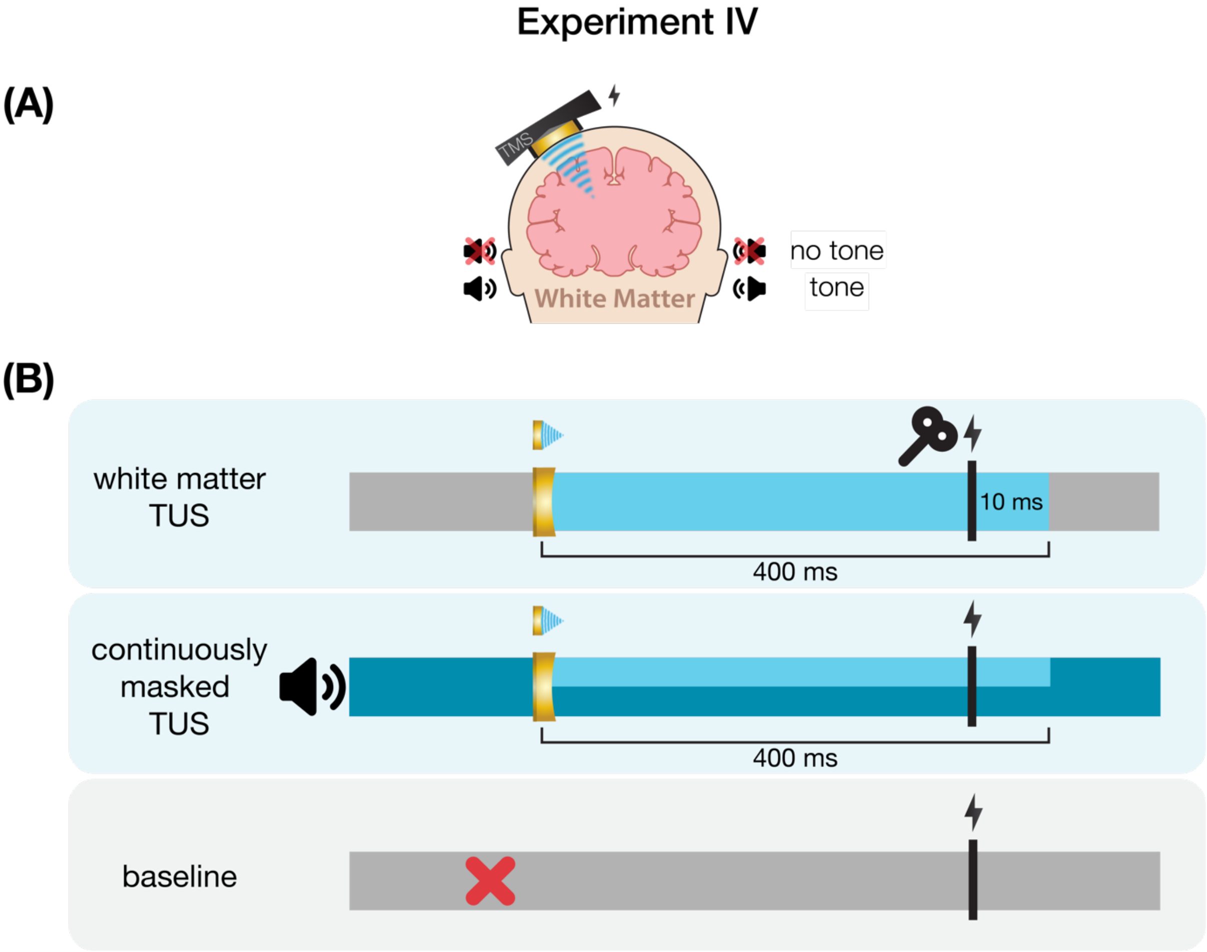

**(A)** Inactive control stimulation of white matter ventromedial to the left-hemispheric hand motor area. **(B)** White matter TUS was administered for 400 ms, with TMS applied 10 ms prior to TUS-offset. In the masked condition, an auditory stimulus was played continuously throughout the entire block.

**Supplementary Figure 4.1 |** Acoustic simulations for Experiment I

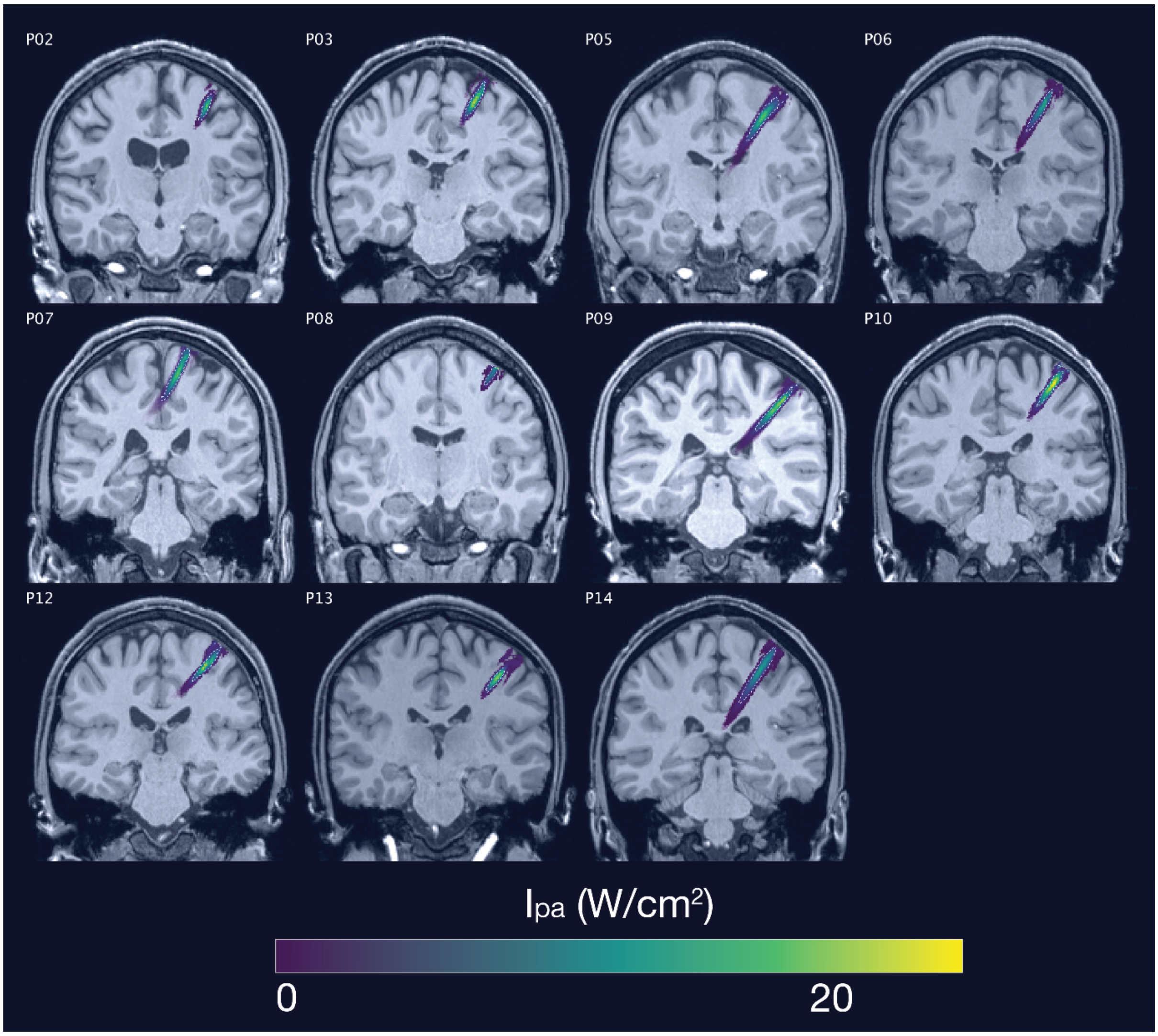

Individual simulations for Experiment I of acoustic wave propagation at 65 W/cm$^2$ free-water stimulation intensity. Simulated pulse-average intensities above 0.15 W/cm$^2$ are depicted. The FWHM of the pressure is indicated by the dashed white line.

**Supplementary Figure 4.2 |** Thermal simulations for Experiment I

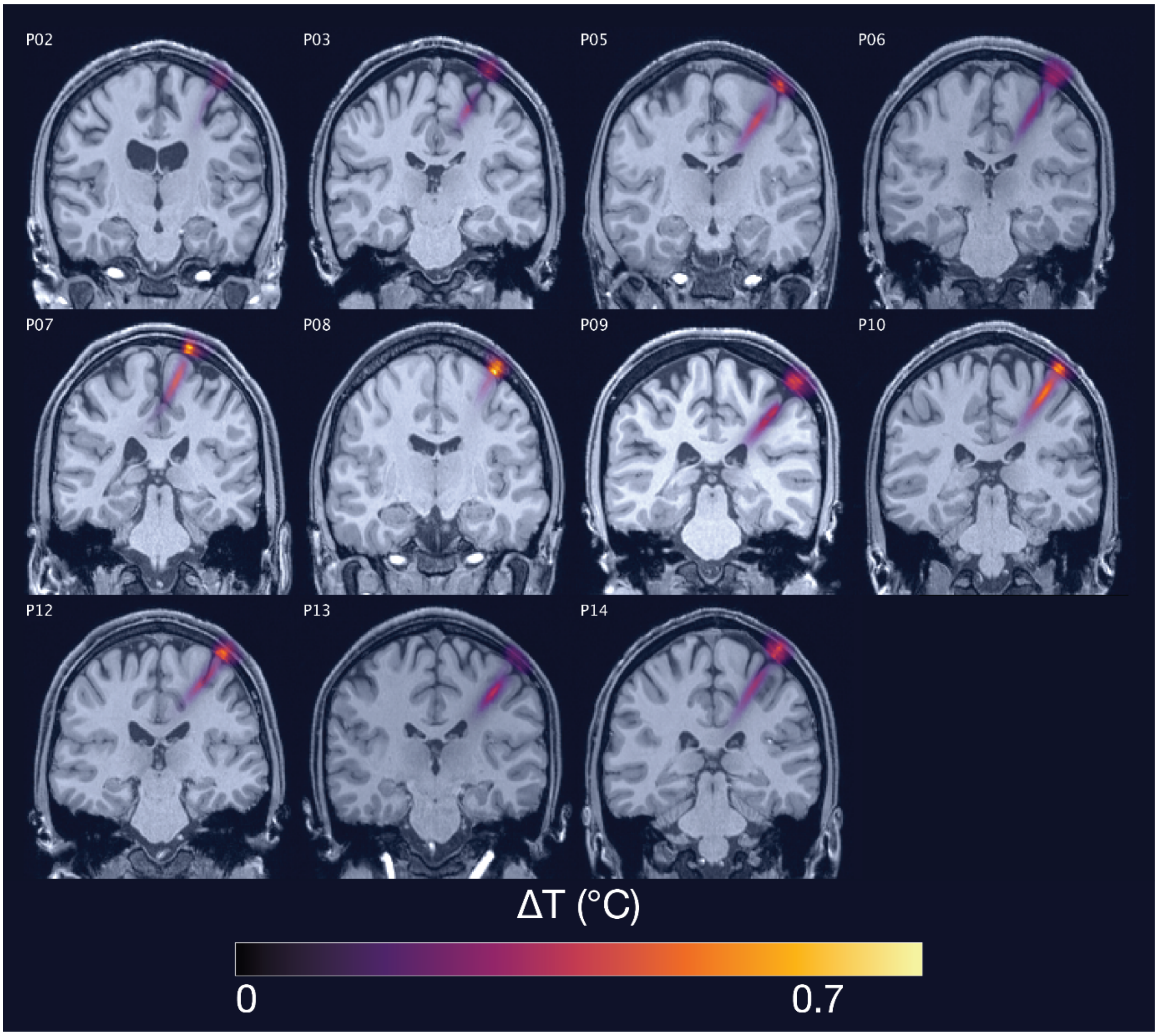

Individual simulations for Experiment I of thermal rise at 65 W/cm$^2$ free-water stimulation intensity.

**Supplementary Figure 4.3 |** Acoustic simulations for Experiment II

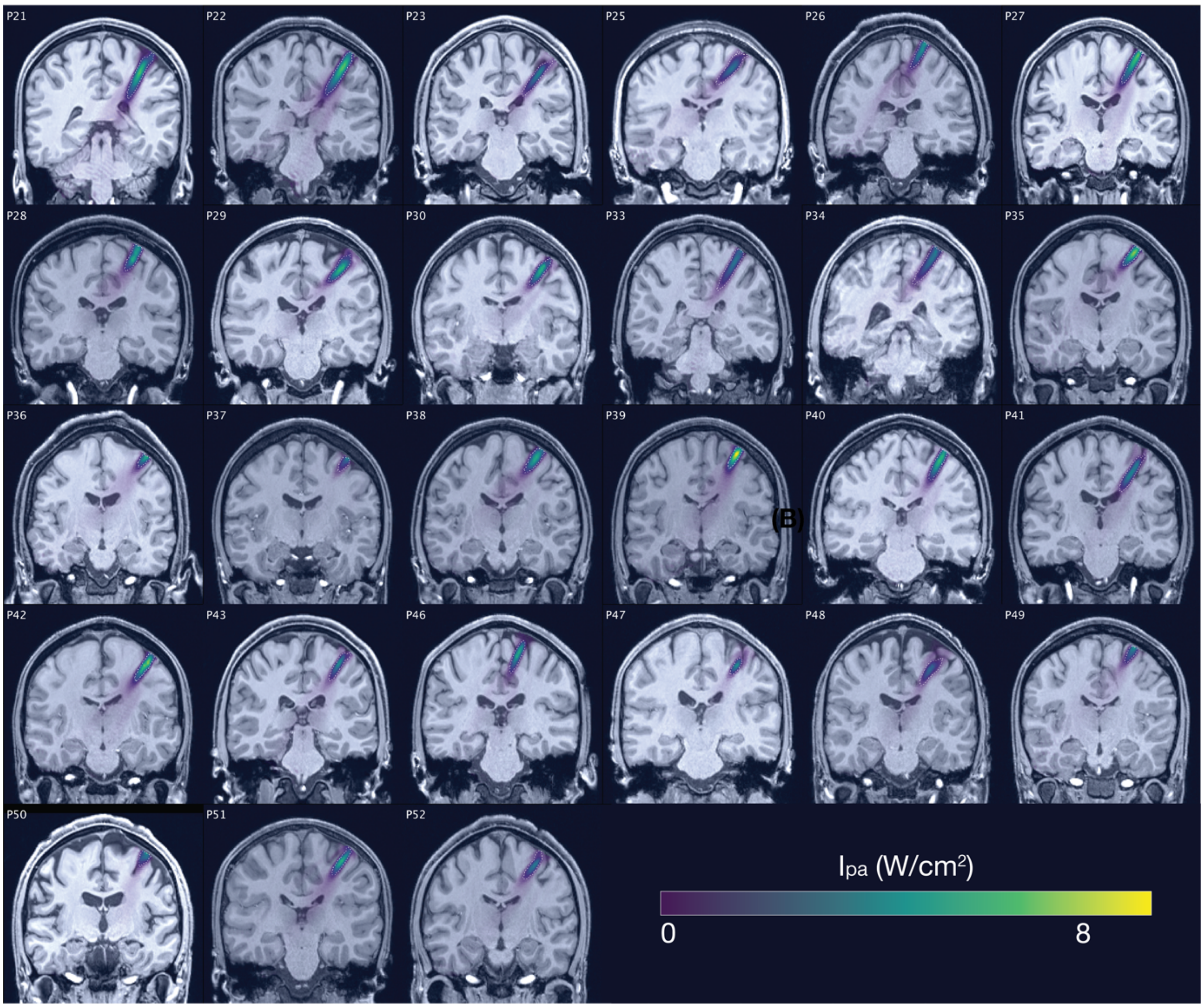

Individual simulations for Experiment II of acoustic wave propagation at 19.05 W/cm$^2$ free-water stimulation intensity. Simulated pulse-average intensities above 0.15 W/cm$^2$ are depicted. The FWHM of the pressure is indicated by the dashed white line.

**Supplementary Figure 4.4 |** Thermal simulations for Experiment II

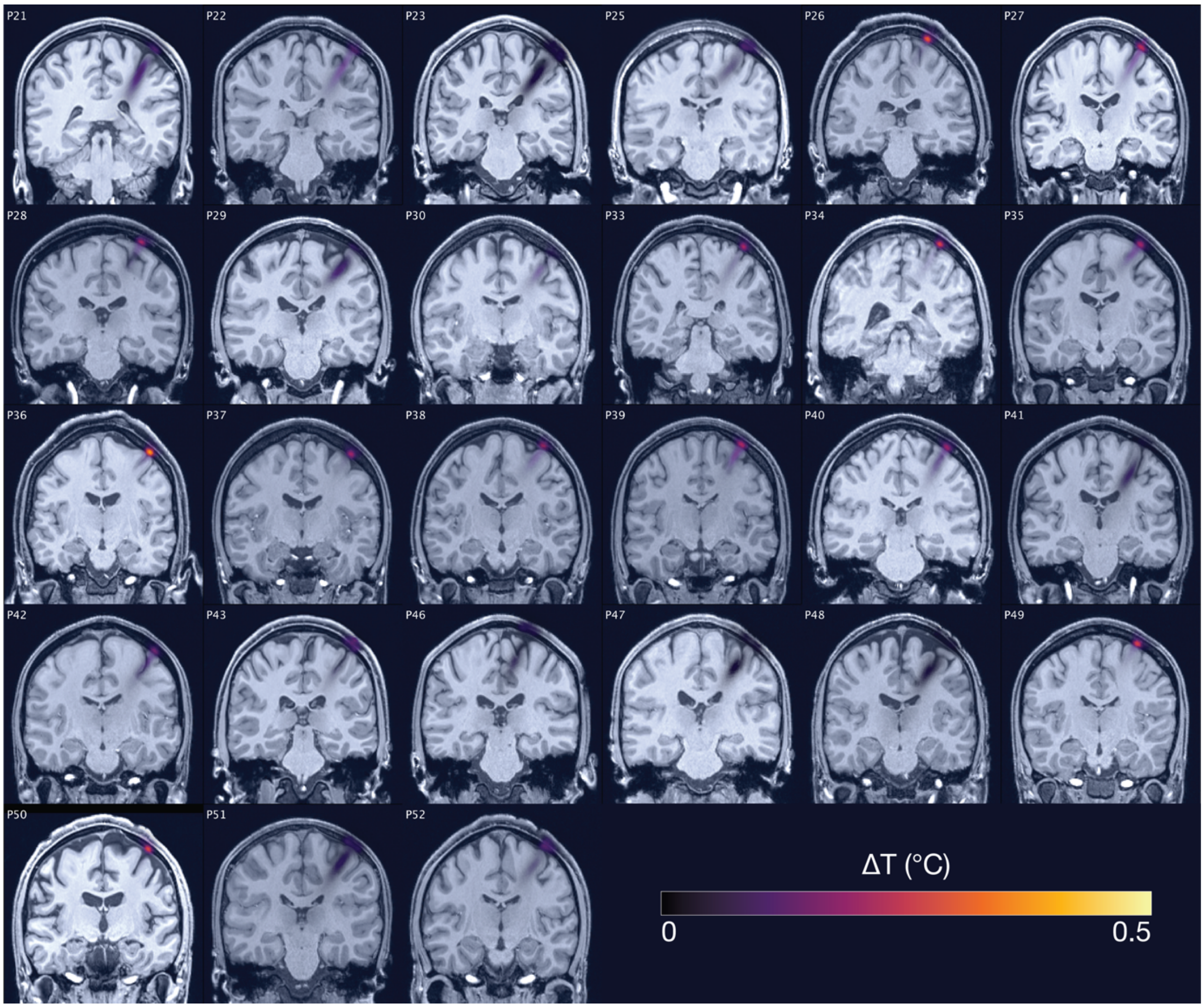

Individual simulations for Experiment II of thermal rise at 19.05 W/cm$^2$ free-water stimulation intensity.

To obtain estimates of acoustic targeting, realized intracranial dosage, and safety indices, we conducted individualized 3D simulations of acoustic and thermal effects for Experiments I and II using the validated k-Wave MATLAB toolbox, a pseudospectral time-domain solver (Treeby & Cox, 2010).

Preprocessing was performed in MATLAB 2019b. Anatomical scans (T1w and T2w) were first segmented into distinct tissues using the SimNIBS headreco tool (Thielscher et al., 2015). We then reoriented the segmented volumes such that the axial plane of the recorded transducer location was parallel to the cartesian axis of the computational grid. Subsequently, the segmented volumes were resampled to an isometric volume size of 0.5 mm and interpolated using the nearest neighbor algorithm. We then extracted the skull, scalp, and brain – including gray and white matter – volumes. To correct minor segmentation errors, the volumes were smoothed by a cubic averaging kernel using a window size of 4 voxels, and re-binarized. The skull layer was additionally combined with a one-voxel outer layer of the filled-in bone volume provided by SimNIBS. Any gaps between skull and skin volumes were assigned to the skull. The skull, scalp, and brain were all included in acoustic and thermal k-Wave simulations, where all voxels not assigned to these tissues were designated as water. Segmented volumes were then cropped to include the whole skull and the transducer, including a perfectly matched layer (PML) of 10 grid points. The points per wavelength given the 0.5 mm grid spacing was 6 for Experiment I ($f$ = 500 kHz) and 12 for Experiment II ($f$ = 250 kHz).

The acoustic properties of sound speed, density, alpha coefficient, and alpha power, as well as the thermal properties of thermal conductivity and heat capacity, were assigned to the skull, scalp, and brain, as reported in ***Table 4*** (Aubry et al., 2022; Hasgall et al., 2022).

We modelled on-target TUS with two-element annular arrays using their respective geometric dimensions and fundamental frequencies for Experiments I and II. The position and orientation of the simulated transducer was extrapolated based on the coordinates of the TMS coil recorded during the experiment with neuronavigation software. We calibrated the simulated source amplitude and phase in free-water to reach the applied free-water stimulation intensities of 32.5 and 65 W/cm$^2$ (Experiment I) and 6.35 and 19.06 W/cm$^2$ (Experiment II). We calibrated the simulated transducer such that the center of the acoustic profile's full-length half-maximum was at the applied depth of 35.1 mm (Experiment I) or 28 mm (Experiment II) from the exit plane of the transducer. The source amplitude and phase resulting in a simulated acoustic profile with the minimum error in relation to the water-tank calibrated acoustic profile provided by the manufacturer were used for acoustic and thermal simulations including the tissue volumes.

Acoustic simulations were then run for three TUS cycles after reaching the steady-state pressure distribution per applied free-water intensity per participant. The maximum pressure map computed by k-Wave was then used to calculate the realized intracranial pulse-average intensity, which was calculated as: $I_{pa} = \frac{p^2}{2c\rho}$ , where *p* is acoustic pressure, *c* is sound speed, and ρ is density. The mechanical index was calculated based on simulated intracranial pressure as: $MI = \frac{p}{\sqrt{f}}$, where *p* is the peak intracranial pressure, and *f* is the fundamental frequency. This formula was also used for Experiments III and IV, but with derated free-water pressures.

The simulated pressure map further served as input for the thermal simulations, where we quantified the maximum thermal rise at each point in the computation grid. Here, simulations were run with a 30% DC for a stimulus duration of 500 ms, corresponding to the applied TUS. To compute a conservative safety estimate of thermal rise, stimulation was split into five 100 ms steps, each with a 30 ms on-period and a 70 ms off-period (actual on-period = 0.3 ms per 1 ms). In addition to thermal rise, the TIC for the highest applied intensity (i.e., 65 W/cm$^2$; pressure = 1.44 MPa) with the highest duty cycle (30%) was calculated as $TIC = \frac{W}{40D}$. The time-averaged acoustic power, W, was calculated as the maximum power (152.34 W) assuming an electrical efficiency of 85% (129.5 W) and considering the 30% duty cycle (129.5 ⋅ 0.3 = 38.85). The aperture diameter, D, was 45 mm. Thus, the highest TIC across all experiments was $\frac{38.85}{40 \cdot 4.5}$ = 0.216.

**Supplementary Figure 5 |** Blinding efficacy (Exp. II).

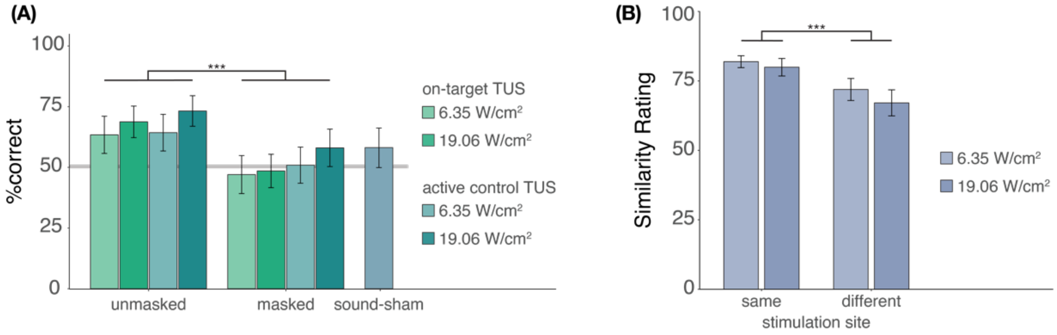

**(A)** An auditory masking stimulus significantly decreased participants' ability to determine whether they had received ultrasonic stimulation. **(B)** However, when masked stimulation was applied over different sites, participants still rated these stimuli as sounding less similar than stimulation applied over the same site. Bars represent means across participants, error bars represent the standard error, and points represent individual participants.

In Experiment II, we investigated whether participants were effectively blinded to ultrasonic stimulation when a time-locked auditory masking stimulus was applied (***Supplementary Figure 5A***). Participants indicated whether they believed TUS was administered across each condition, excluding baseline. Participants completed 4 trials per condition, resulting in 36 trials in total, presented in pseudorandomized order where each condition is presented randomly in sets of 9 trials. We used a mixed effects logistic regression to predict participant response (stimulation yes/no) as a function of stimulation site (on-target/active control), free-water intensity (6.35 and 19.05 W/cm$^2$), and masking (no mask/masked) with random intercepts by participant. This test revealed significantly lower detection rates when masking was applied ($b$ = -0.90, $SE$ = 0.31, $z$ = -2.87, $p$ = 0.004), but showed no significant effect of stimulation site ($b$ = 0.05, $SE$ = 0.31, $z$ = 0.16, $p$ = 0.875) or intensity ($b$ = 0.30, $SE$ = 0.31, $z$ = 0.95, $p$ = 0.342), nor any significant interactions (all $p$ > 0.6). Only ultrasonic stimulation applied at higher intensities (19.06 W/cm$^2$) in the absence of an auditory masking stimulus were detected by participants at an above-chance rate (i.e., 50%; on-target: $Z$ = 2.6, $p$ = 0.011; active control: $Z$ = 2.9, $p$ = 0.004). Detection rates for lower intensities, masked stimulation, and sound-sham conditions did not differ significantly from chance, as revealed by post-hoc one-sample Wilcoxon signed rank tests (all $Z$ < 1.7, $p$ > 0.09). Taken together, these results show that the masking used in Experiment II successfully reduced participants' ability to determine whether they had received stimulation.

It is possible that, while blinding to TUS versus no-TUS is successful for explicit judgements, slight audibility differences between on-target and active control stimulation persist even when stimulation is masked. To test for audible differences between stimulation sites (***Supplementary Figure 5B***), participants were exposed to two subsequent masked TUS trials. These two stimuli were either the same (i.e., [on-target + on-target] or [active control + active control]) or different (i.e., [on-target + active-control/active-control + on-target]). These two trials were presented at both intensities (6.35 and 19.05 W/cm$^2$), to account for the louder auditory confound associated with higher intensity stimulation. Following each pair of trials, participants were asked to rate their similarity on a visual analog scale (0 = very dissimilar, 100 = very similar) across 8 trials in total. We conducted a two-way repeated measures ANOVA predicting similarity rating by stimulation site (same/different) and stimulation intensity (6.35/19.05 W/cm$^2$). A two-way repeated measures ANOVA revealed that masked stimulation over the two different sites sounded less similar than masked stimulation over the same site (***Supplementary Figure 5B***; $F(1,26)$ = 14.00, $p$ = $9{\cdot}10^{-4}$) while no significant main effect of, or interaction with, intensity was observed (main effect: $F(1,26)$ = 1.98, $p$ = 0.171; interaction: $F(1,26)$ = 0.26, $p$ = 0.613). These results suggest that, while masking may effectively blind participants, thus reducing detection of ultrasonic stimulation, slight audible differences between different stimulation sites may still persist.

**Supplementary Figure 6 |** No evidence for direct intracranial dose-response effects (Exp. II)

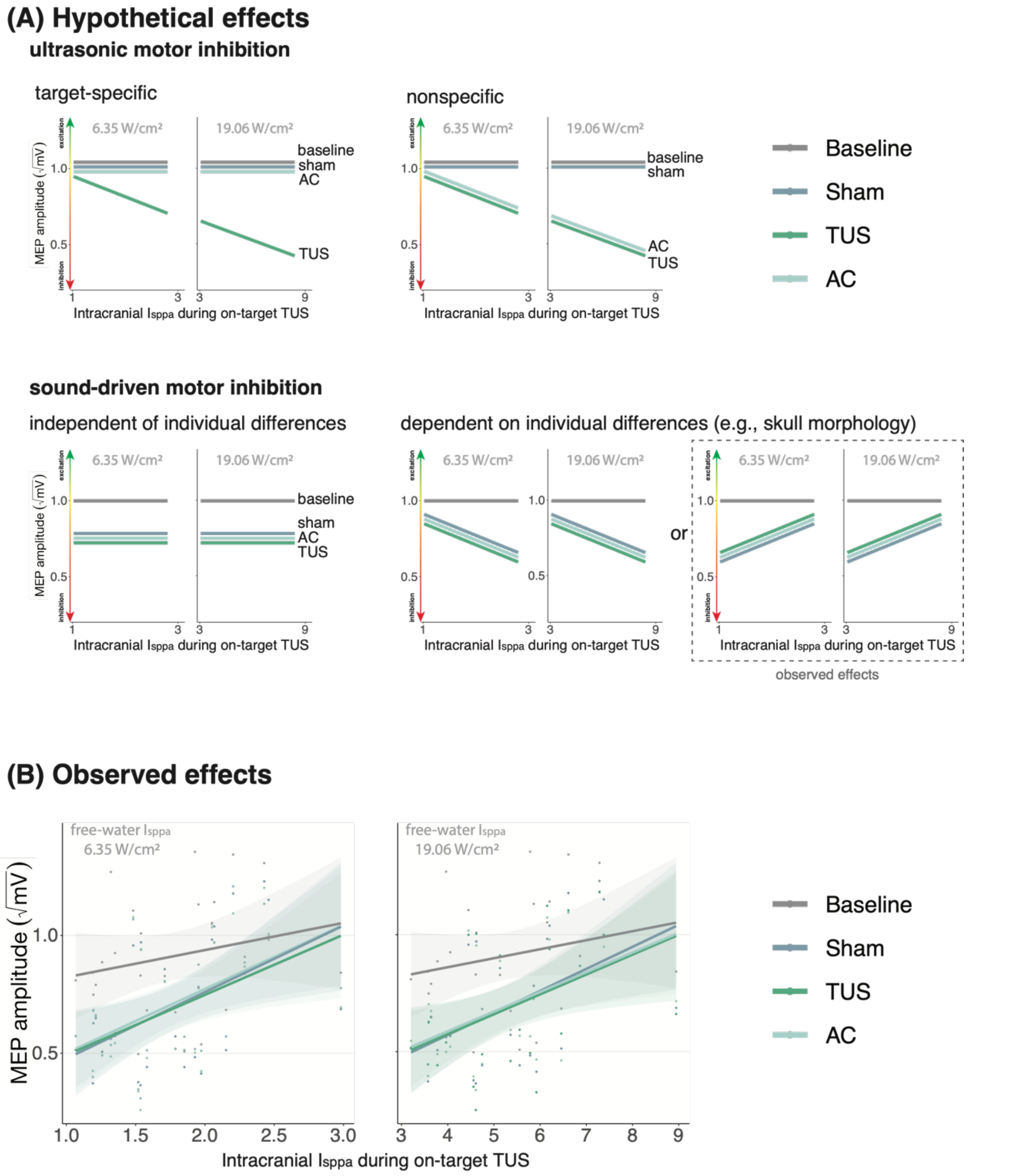

**(A)** Hypothetical effects. A target specific dose-response effect of TUS would be reflected by a change in MEP amplitude with increasing intracranial intensity only for on-target TUS (top left), whereas a nonspecific effect of TUS would show the same 'dose-response' for both on-target and active control conditions (top right). If there is sound-driven inhibition, a motor inhibitory effect would be observed for on-target, active control, and sound-only sham (bottom left). If sound-driven inhibition would be subject to individual differences, for example in skull morphology which correlates with intracranial intensity, there would be a correlation between MEP amplitude and intracranial intensity that exists for on-target, active control, and sound-sham conditions (bottom right). The latter corresponds with the observed effects. **(B)** Average square root corrected MEP amplitude, plotted separately for baseline, sham, TUS, and active control conditions, across simulated intracranial intensities. For each participant, we estimated the intracranial ultrasound intensity at the hand M1 target, represented on the x-axis, and the average MEP amplitude for all four conditions. Results for on-target and active control delivered at 6.35 W/cm$^2$ free-water $I_{sppa}$ are depicted on the left, and at 19.06 W/cm$^2$ free-water $I_{sppa}$ are depicted on the right. Please note that, for reference, the baseline and sham conditions are duplicated across both plots. There is no significant difference between on-target TUS and control conditions across inter-individual intracranial intensities. Points represent the average MEP amplitude per participant per condition. The shaded area represents the 95% CI.

In Experiments I & II, we tested potential dose-response effects of TUS by applying stimulation at multiple free-water intensities. Here, no significant effect of administered intensity was observed. However, the efficacy of TUS likely depends on realized intracranial intensities. Therefore, we ran 3D acoustic simulations to estimate the intracranial intensity during on-target TUS. It was not appropriate to combine data from Experiments I and II given the different fundamental frequencies and stimulation depths applied.

Should direct and spatially specific neuromodulation be taking place, we would expect to see a dose-response effect where MEP amplitude changes with intracranial intensity during on-target TUS, and not during control conditions. Therefore, the critical test to provide evidence of direct neuromodulation is a comparison of the dose-response relationship between on-target TUS and control conditions. To this end, we ran simple linear models for Experiment II, which had a sufficient sample size (n = 27) to assess inter-individual variability. We found no significant difference in the TUS-MEP relationship between on-target TUS and control conditions (active control free-water 6.35 W/cm$^2$: $b$ = -1.02, $SE$ = 3.66, $t(24)$ = -0.28, $p$ = 0.783; active control free-water 19.06 W/cm$^2$: $b$ = 0.04, $SE$ = 1.66, $t(24)$ = 0.02, $p$ = 0.983; sound-sham free-water 6.35 W/cm$^2$: $b$ = 0.58, $SE$ = 4.78, $t(24)$ = 0.12, $p$ = 0.90; sound-sham free-water 19.06 W/cm$^2$: $b$ = -0.66, $SE$ = 1.92, $t(24)$ = -0.35, $p$ = 0.733). We conclude that there is no evidence for a direct neuromodulatory intracranial dose-response relationship. This interpretation is in line with our findings when testing free-water dose-response effects of TUS.

Notably, given that for all individuals the same ultrasound intensity was applied at the source outside the skull, any intracranial differences in intensity are primarily driven by individual skull characteristics, such as skull thickness. These individual characteristics are expected to co-vary with TMS parameters, such as the TMS intensity (%MSO) required to obtain a 1mV MEP. It is conceivable that such co-varying relationships can drive overall effects of MEP amplitude and MEP inhibition. Indeed, we observe a significant effect of intracranial intensity on the difference in MEP amplitude between on-target TUS at 6.35 W/cm$^2$ and baseline ($b$ = 19.46, $SE$ = 7.91, $t(24)$ = 2.46, $p$ = 0.022), as well as a trend for on-target TUS at 19.06 W/cm$^2$ ($b$ = 5.77, $SE$ = 3.01, $t(24)$ = 1.92, $p$ = 0.067). Importantly, this is also observed for the sound-sham condition ($b$ = 19.65, $SE$ = 7.58, $t(24)$ = 2.59, $p$ = 0.016), where direct neuromodulation is impossible. These observations emphasize the need for rigorous control conditions to support strong inferences of TUS neuromodulation. In summary, in this experiment, no evidence for a specific and direct neuromodulatory dose-response effect was observed.

**Supplementary Figure 7 |** Temporal dynamics (Exp. II).

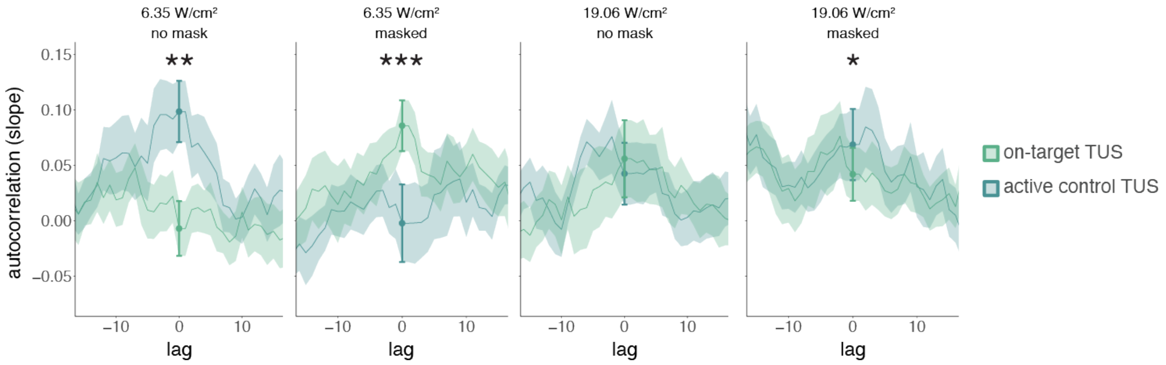

We operationalized autocorrelation as the slope between MEP amplitude on a given trial (*t*) and its preceding baseline MEP amplitude (*t-1*) for each condition. Statistical inference is drawn at the current trial (lag = 0). Dots and error bars reflect the mean and standard error of the slopes across participants for each level of on-target and active control TUS. Additionally, for visualization purposes we display autocorrelation for shifts in the preceding baseline timeseries, with lags ranging from -15 to +15 trials. The line depicts the mean slope across participants for each lag, and the shaded area represents the standard error. * $p < .05$, ** $p < 0.01$, *** $p < 0.001$.

While studies often focus on quantifying excitatory or inhibitory effects of TUS, it can be informative to additionally examine how TUS affects neural dynamics in the time domain. Here, we conduct a preliminary exploration of whether autocorrelative measures between a given trial (*t*) and its preceding measure of baseline motor cortical excitability (*t-1*) yield insight into possible introduction of noise by TUS. To this end, we ran a linear mixed model predicting square root corrected MEP amplitude by 'previous baseline' amplitude, as well as 'stimulation site' (on-target/active control), 'intensity' (6.35/19.06 W/cm$^2$), and 'masking' (no mask/masked). Previous baseline MEP amplitudes were mean centered and standardized on a participant level. Random intercepts and slopes were included for 'stimulation site', 'intensity', 'previous baseline', and their interaction, permitting model convergence. This test revealed a significant four-way interaction ($F(1,5222) = 26.10$, $p = 3{\cdot}10^{-7}$, $\eta_p^2 = 5{\cdot}10^{-3}$) and an accompanying main effect of previous baseline MEP amplitude ($F(1,26) = 5.70$, $p = 0.024$, $\eta_p^2 = 0.18$). Follow-up LMMs for unmasked and masked stimulation separately both revealed a significant three-way interaction between 'stimulation site', 'intensity', and 'previous baseline' (unmasked: $F(1,176) = 7.32$, $p = 0.007$, $\eta_p^2 = 0.04$; masked: $F(1,30) = 12.10$, $p = 0.002$, $\eta_p^2 = 0.28$). Further follow-up LMMs for each level of masking and intensity were used to test the effects of 'stimulation site', 'previous baseline', and their interaction. These LMMs revealed a significant interaction for all but unmasked 19.06 W/cm$^2$ stimulation (**Supplementary Fig. 7**; no mask 6.35 W/cm$^2$: $F(1,1293) = 8.36$, $p = 0.004$, $\eta_p^2 = 6{\cdot}10^{-3}$; no mask 19.06 W/cm$^2$: $F(1,1283) = 1.13$, $p = 0.288$, $\eta_p^2 = 9{\cdot}10^{-4}$; masked 6.35 W/cm$^2$: $F(1,1287) = 13.43$, $p = 3{\cdot}10^{-4}$, $\eta_p^2 = 0.01$; masked 19.06 W/cm$^2$: $F(1,1282) = 5.76$, $p = 0.017$, $\eta_p^2 = 4{\cdot}10^{-3}$).

**Supplementary Fig. 7** shows the temporal autocorrelation, as indexed by the linear regression slopes of the relationship between MEP amplitude on the previous baseline trial and the current test trial. Statistical inference is drawn at the current trial (zero-lag, indicated with the filled dots and error bars). For visualization only, the autocorrelation is calculated for a range of time lags, shifting the timeseries of preceding baseline excitability from 15 trials in the past to 15 trials in the future. This shows, as expected, that autocorrelation and the modulation thereof disappear at larger absolute time lags. Figure 4D in the main manuscript shows the difference in autocorrelation between on-target and active control TUS (**Fig. 4D**; e.g., masked 6.35 W/cm$^2$ on-target TUS slope – masked 6.35 W/cm$^2$ active control slope). For sound-sham and baseline, the difference score was calculated based on the mean over all active control trials.

In sum, these preliminary exploratory analyses could point towards TUS introducing temporally specific neural noise to ongoing neural dynamics in a dose-dependent manner, rather than simply shifting the overall excitation-inhibition balance. One possible explanation for the discrepancy between trials with and without auditory masking is the difference in auditory confound perception, where without masking the confound's volume differs between intensities, while with masking this difference is minimized. Future studies might consider designing experiments such that temporal dynamics of ultrasonic neuromodulation can be captured more robustly, allowing for quantification of possible state-dependent or nondirectional perturbation effects of stimulation.

**Supplementary Figure 8 |** Subjective report of TUS audibility (Exp. IV).

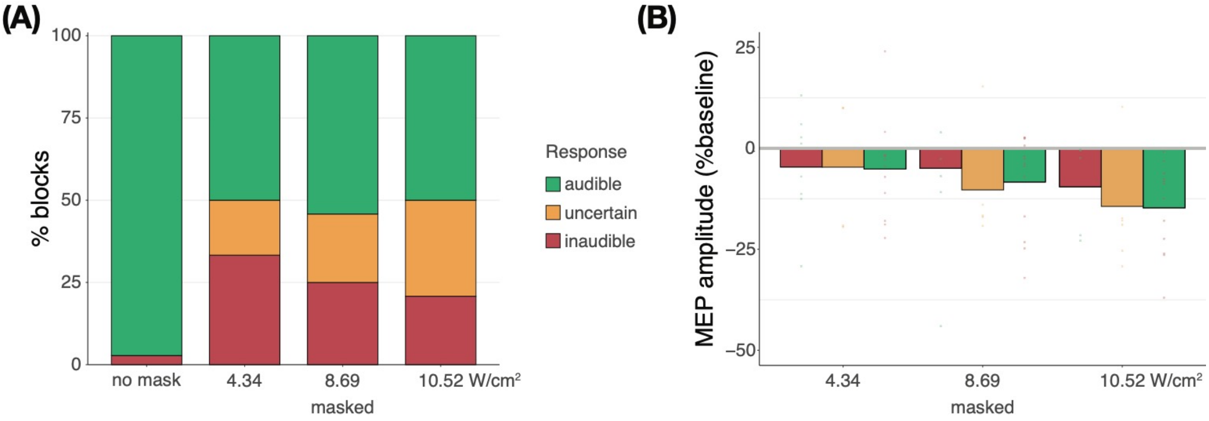

**(A)** Depicting the percentage of blocks for which participants reported a condition as audible, uncertain, or inaudible. When no mask was applied, TUS was audible (green) in nearly all cases. During continuous masking, TUS was experienced as inaudible (red) less often at higher stimulation intensities. Descriptively, reports of inaudibility scale with MEP amplitudes such that in conditions where TUS was inaudible more frequently, less motor inhibition was observed. **(B)** Motor inhibitory effects during masked stimulation, split by audibility. Descriptively, when masking rendered TUS inaudible, less motor inhibition was observed than when participants were uncertain or when stimulation was heard.

We propose that the results of Experiment IV reflect the impact of auditory confound volume. Here, the volume of the confound can be regarded as the salience of a cue for the upcoming TMS pulse. When masking is applied, the salience of this cue is diminished. Correspondingly, less motor inhibition was observed when stimulation was masked, as opposed to when no masking stimulus was administered and the auditory confound was clearly audible.

During continuous masking, more motor inhibition was observed at higher auditory confound volumes (i.e., intensities). This is likely because the confound can then be perceived, albeit potentially unconsciously. Indeed, in Supplementary Figure 4B, we also see that participants who rated masked stimulation as uncertain or audible demonstrated more inhibition. Taken together, we suggest that 'dose-response' effects of auditory confound volume are being observed. As the confound approaches and exceeds the boundary of (subconscious) audibility, sufficiently salient cueing of the upcoming TMS pulse takes place, thus evoking proportional motor inhibitory effects.
